# Chimeric Collagen-like Proteins with Tunable Structural Heterogeneity for Precise Control and Targeted Doxorubicin Delivery

**DOI:** 10.1101/2025.03.09.641987

**Authors:** Yan Xia, Yimiao Li, Jinxia Huang, Jian Feng, Jiaqi Li, Chuan Liu, Shuang Jia, Shad Man, Shubin Li, Rile Wu, Zhiyan Ren, Yukun Zang, Liping Wang, Xing Liu, Jie Wang, Yinan Sun, Liyao Wang, Xinyu Li

## Abstract

Protein-based drug delivery systems offer advantages such as genetic tunability, structural homogeneity, and high biocompatibility. However, precisely controlling the assembly of protein nanoparticles and establishing correlations between their macroscopic properties and molecular architecture remain significant challenges. Here, inspired by chimeric collagen-like proteins, we rationally designed a series of de novo chimeric proteins comprising an N-terminal globular domain fused to a linear collagen-like domain. These structurally heterogeneous proteins were conjugated with doxorubicin (DOX) and homogenized to form chimeric collagen-like protein–DOX nanocomplexes (CDCNs). By fine-tuning the protein structure and domain organization, we achieved precise control over CDCN architecture and drug release kinetics. Furthermore, to enhance tumor specificity, triple-negative breast cancer (TNBC)- and glioblastoma (GBM)-targeting peptides were genetically fused to the chimeric proteins, respectively. In both in vitro and in vivo models of TNBC and GBM, CDCNs facilitated selective tumor accumulation, enhanced cellular uptake, and promoted apoptosis while minimizing off-target toxicity. This work establishes a strategy for designing protein-based nanoplatforms with programmable structures and tunable functionalities, offering promising potential for precisely controlling of protein-based delivery system.

## Introduction

Proteins, as fundamental biomolecules, are characterized by homogeneity, biocompatibility, and modularity, making them ideal for engineering nanoscale drug delivery systems^1-4^. Through biochemical and genetic modifications, proteins can be tailored for controlled drug release and targeted delivery^4, 5^. However, achieving precise control over protein nanoparticle structures and correlating their macroscopic properties with underlying molecular architecture remains a challenge^6^.

Commonly used globular proteins for nanocarrier applications include bovine serum albumin (BSA)^7-9^ and human serum albumin (HSA)^10-12^. Nanoparticles from these proteins are typically fabricated via emulsification or desolvation, where desolvating agents induce protein aggregation^1, 3, 11, 13^. Although these methods are simple and reproducible, issues such as poor homogeneity, low stability, and variable drug encapsulation efficiency limit their application^1, 3^. In contrast, linear protein-based nanocarriers, such as elastin^14-16^ and collagen^4, 17, 18^, offer advantages in structural flexibility and tunability^16, 19^. These carriers are formed by conjugating therapeutic agents to the protein backbone, followed by self-assembly driven by hydrophobic interactions^14-16^. However, the resulting wool-like structures may reduce targeting efficiency due to the compromised spatial arrangement of modified targeting peptides, affecting delivery specificity^20^.

Due to the limitations of both globular and linear proteins as delivery systems, combining linear proteins with rigid structures that integrate the advantages of both offers an optimal strategy for designing protein-based carriers^14, 20^. For example, fusing a silica-binding sequence to the C-terminus of linear elastin sequences, followed by coordination with silica nanoparticles, results in hybrid protein carriers that significantly enhance drug release control and targeting ability compared to native elastin proteins^14^.

Recently, recombinantly synthesized bacterial collagen-like proteins (CLPs) have garnered significant attention for their potential applications in tissue engineering^21-23^. These CLPs consist of a globular domain followed by a linear collagen-like domain, resemble mammalian collagen and contain the characteristic (Gly-Xaa-Yaa)_n_ sequence.^23-25^. CLPs lack bioactive sequences, providing a customizable platform for integrating biologically relevant motifs^25, 26^. Despite their promise, the potential of CLPs as drug carriers remains underexplored.

Inspired by the chimeric structural characteristics of CLPs, this study introduces chimeric CLP-like proteins with an N-terminal globular domain, a central linear domain, and C-terminal conjugation sites^15^. These chimeric structures were precisely engineered through de novo design and genetic encoding of these three key domains. By exploiting the inherent structural heterogeneity of CLPs, doxorubicin (DOX) was conjugated at the C-terminal site to form chimeric CLP-DOX conjugated nanoparticles (CDCNs)^15, 20^. The structural diversity of CDCNs was analyzed for drug release profiles, structural dynamics, and anti-tumor efficacy. To further enhance specificity, targeting peptides against triple-negative breast cancer (TNBC)^27^ and glioblastoma (GBM)^28^ were genetically encoded into the chimeric proteins. The in vitro and in vivo anti-tumor efficacy of the CDCNs was evaluated, demonstrating their potential as targeted protein-based drug delivery systems. Overall, this study presents a novel strategy for designing chimeric CLP-like proteins as versatile drug carriers. By offering precise control over nanoparticle structure and drug release properties, this approach has potential applications in cancer therapy and beyond.

## Results

### Design strategy and preparation of CDCNs

Building upon the structural principles of CLPs, we designed a series of chimeric proteins derived from *Streptococcus pyogenes* Scl2 protein^21^. A rigid globular domain was positioned at the N-terminus to enhance stability (Fig. 1B), followed by a collagen-like sequence with repetitive (Gly-Xaa-Yaa)_n_ motifs, enabling the formation of a coiled-coil helical structure (Fig. 1B)^29^. The C-terminus incorporated repetitive (GC)_n_ sequences to serve as conjugation sites for doxorubicin (DOX) attachment (Fig. 1B)^15^. DOX was first activated via N-β-maleimidopropionic acid hydrazide (BMPH) to generate a maleimide-functionalized form. This activated DOX was then covalently conjugated to cysteine residues within the CLP-like proteins using a Michael addition reaction (Fig. 1A)^14, 15, 20^. The resulting chimeric protein-DOX conjugates (CDCNs) were prepared through homogenization-induced micellization (Fig. 1C)^14^. Based on the self-assembly properties of these protein nanoparticles, we hypothesize that the flexible, DOX-conjugated linear domains are encapsulated in the core, while the rigid globular domains form the nanoparticle shell (Fig. 1D)^14, 16^. As a result, key macroscopic characteristics of the CDCNs—such as size, morphology, and drug release profiles—are governed by the length of the linear domain, the extent of DOX conjugation, and the rigidity of the globular domain (Fig. 1D).

**Figure 1.**
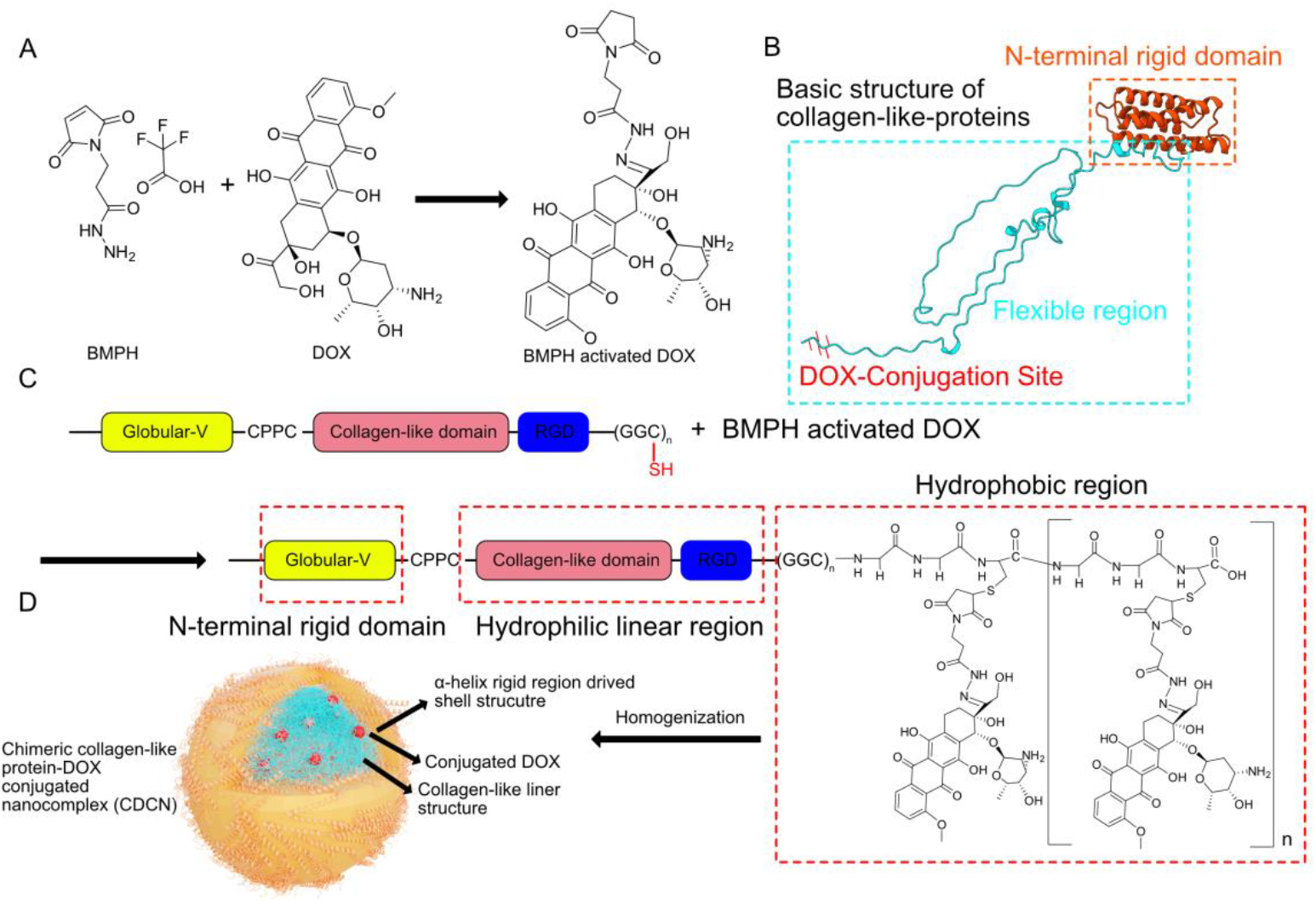
Design and preparation mechanism of CDCNs. A. Schematic representation of DOX activation. DOX was functionalized with BMPH, introducing a terminal maleimide group. B. Structural characteristics of the chimeric protein. The protein consists of three main domains: an N-terminal rigid globular domain, a flexible linear structure, and C-terminal conjugation sites for DOX. C. Reaction scheme for the conjugation of activated DOX to the protein carrier. The maleimide group of the activated DOX reacts with the thiol group of cysteine residues in the protein via a Michael addition reaction. D. Proposed macrostructure of CDCNs. The DOX-conjugated linear domains are encapsulated within the core, while the rigid globular domains form the nanoparticle shell. CDCNs are formed through homogenization-induced micellization of the conjugated proteins.

### The Length of the Linear Structure Determines the Size and Release Rate of CDCNs

To explore the relationship between the microstructure of protein carriers and the macrostructure of their corresponding CDCNs, we constructed a series of chimeric CLPs with varying lengths of the linear domain (Fig. 2A, Fig. S1). Based on the structure of Scl2, which includes 22 repeats of the (Gly-Xaa-Yaa) sequence, we designed three constructs with 22, 44, and 66 repeats, denoted as V_α222_CLP1, V_α222_CLP2, and V_α222_CLP3 (Fig. 2A, Fig. S1)^29^. We found that CLPs failed to fold correctly without the inclusion of an N-terminal globular domain, highlighting its essential role in protein stability. To address this, we incorporated the rigid α-helix Scl2 globular domain (222 base pairs) at the N-terminus of each construct (Fig. 2A, Fig. S1), enabling efficient expression and purification of the chimeric proteins.

**Figure 2.**
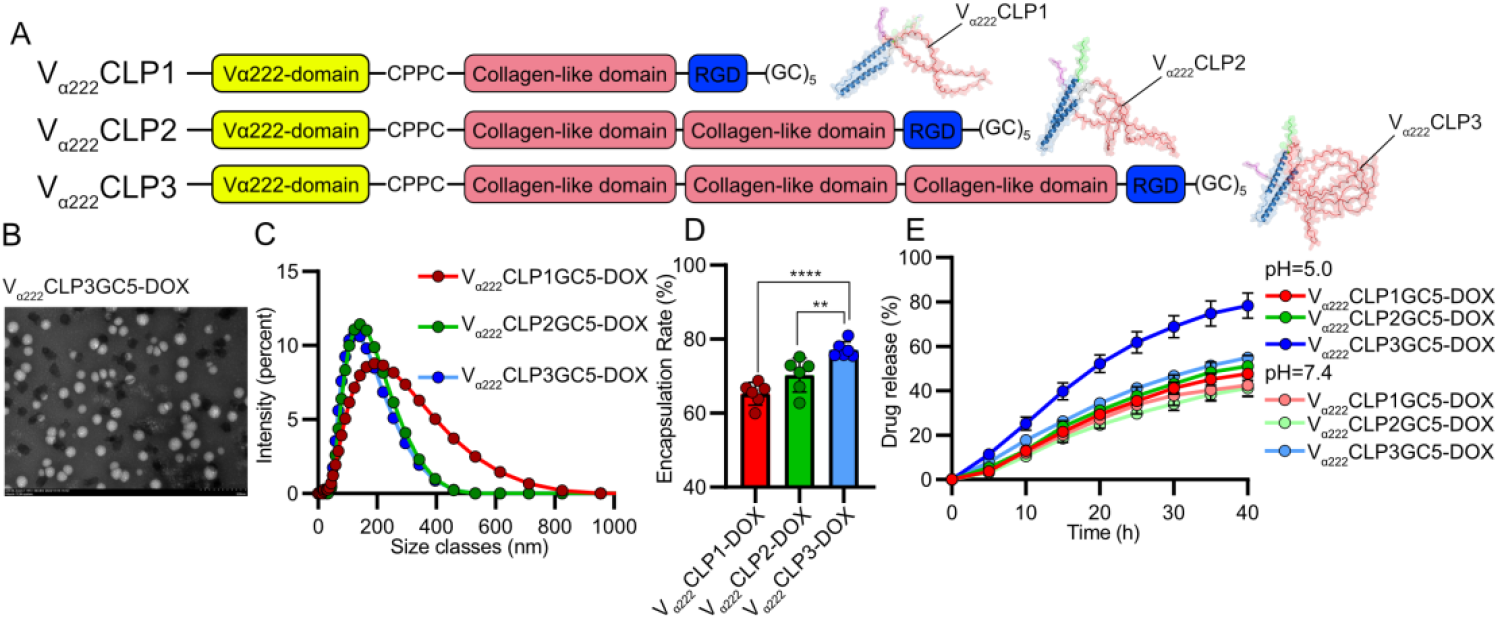
Formation of CDCNs with Different Linear Structure Lengths. A. Structural design and predicted structure of CLPs. The structure of CLPs with varying lengths of linear domains was predicted using AlphaFold 3. The linear domain is shown in red, the rigid α-helix domain in blue, and the conjugation site for DOX, consisting of the (GC)^5^ sequence, is highlighted in purple. B. TEM image of V_α222_CLP3-DOX CDCNs. TEM image shows the homogeneous spherical morphology of the CDCNs formed from V_α222_CLP3-DOX conjugates. C. Size distribution of CDCNs formed from CLPs with different linear structure lengths. DLS analysis of CDCNs formed from CLPs reveals the impact of linear domain length on nanoparticle size. D. Encapsulation efficiency of CDCNs. Encapsulation efficiency of DOX within CDCNs formed from CLPs of varying linear domain lengths was assessed, showing differences in the ability to encapsulate DOX. E. DOX release profiles from CDCNs. The release rate of DOX from CDCNs formed from CLPs with varying linear structure lengths was examined at pH 5 and pH 7, demonstrating the effect of linear domain length on drug release kinetics in different pH environments.

Following the design and purification of these CLPs, activated DOX was conjugated via a maleimide-thiol reaction, and the resulting conjugates were processed through homogenization to form CDCNs (Fig. 1C). Transmission electron microscopy (TEM) images confirmed the successful formation of homogeneously spherical nanoparticles (Fig. 2B). Interestingly, each doubling of the linear domain length led to a reduction in nanoparticle size, suggesting that the increased hydrophilicity of the longer linear sequences promotes aggregation of hydrophobic DOX molecules, resulting in a denser internal structure. This hypothesis is supported by the finding that V_α222_CLP3-DOX CDCNs, which contain the longest linear sequence, exhibited the highest drug encapsulation efficiency, likely due to enhanced aggregation of the DOX payload (Fig. 2D).

The DOX release rate from CDCNs exhibited a proportional relationship with the length of the linear domain. V_α222_CLP3-DOX CDCNs released approximately 50% of their DOX content within 20 hours, while V_α222_CLP2-DOX CDCNs required about 40 hours to reach the same level of release (Fig. 2E). This pattern of accelerated release in CDCNs with longer linear domains was consistent under both pH 5 and pH 7 conditions (Fig. 2E). The increased release rate is likely attributed to the higher hydrophilicity and flexibility of the longer linear sequences, which may facilitate nanoparticle swelling in physiological environments, thereby promoting faster drug release.

### The Opposite Effects of α-Helix and β-Sheet Rigid Domains on CDCNs

To explore the impact of the rigid domain structure on the properties of CDCNs, we engineered chimeric proteins with different N-terminal globular domains. We first used the Scl2 protein’s α-helix domain (222 base pairs)^30^, and designed additional constructs: a single α-helix (102 base pairs) and a pair of reverse-paired α-helices (666 base pairs) (Fig. 3A, D, Fig. S2A). These three different chimeric proteins (V_α102_CLP3, V_α222_CLP3, and V_α666_CLP3) were used to construct CDCNs under identical conditions (Fig. 3A, D, Fig. S2A).

**Figure 3.**
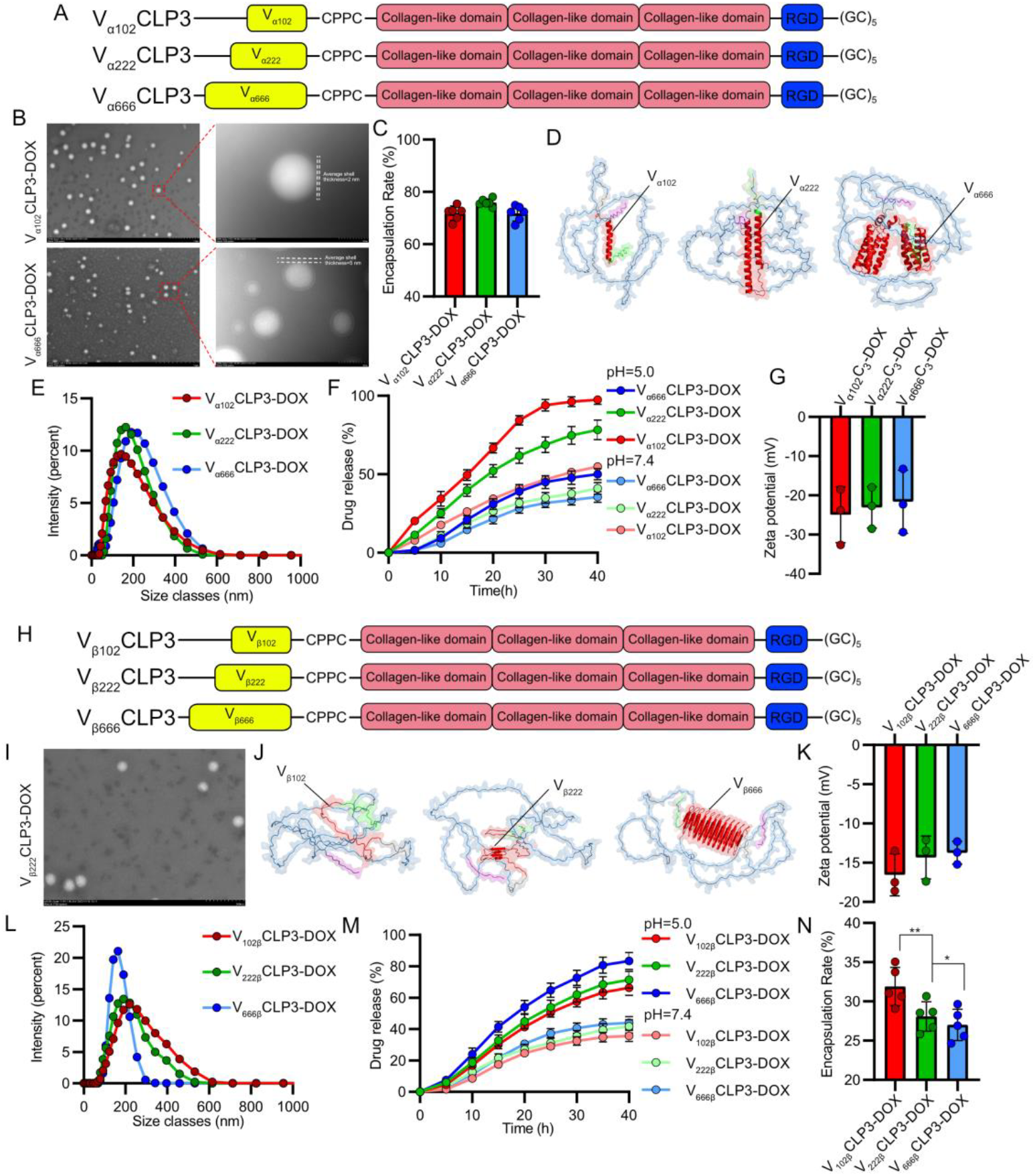
Effects of N-terminal globular structure heterogeneity on CDCNs. A. Schematic representation of the design of N-terminal α-helix heterogeneity in chimeric CLPs. The CLPs were engineered to contain a heterogeneous N-terminal α-helix domain and 66 repeats of the (Gly-Xaa-Yaa) sequence. B. TEM images illustrating the macrostructure of V_α102_CLP3-DOX and V_α666_CLP3-DOX nanoparticles. C. Encapsulation efficiency of CDCNs. The heterogeneity of the N-terminal α-helix domain had minimal effect on the encapsulation efficiency. D. Protein structures were predicted using AlphaFold 3, with the linear domain depicted in blue, the rigid α-helix domain in red, and the DOX conjugation site in purple. E. Size distribution of CDCNs derived from α-helix heterogeneity CLPs. The nanoparticle size increased slightly with the complexity of the N-terminal α-helix domain. F. Drug release profiles for CDCNs derived from α-helix heterogeneity CLPs. The release rate was modulated by the complexity of the N-terminal α-helix domain. G. Zeta potential of CDCNs derived from α-helix heterogeneity CLPs. H. Schematic representation of the design of N-terminal β-sheet heterogeneity in chimeric CLPs. These CLPs were engineered to contain a heterogeneous N-terminal β-sheet domain and 66 repeats of the (Gly-Xaa-Yaa) sequence. I. TEM images of V_β222_CLP3-DOX, showing minimal effect of β-sheet domain heterogeneity on the macrostructure of the CDCNs. J. The protein structure was predicted by AlphaFold 3, with the linear domain shown in blue, the rigid β-sheet domain in red, and the DOX conjugation site in purple. K. Zeta potential of CDCNs derived from β-sheet heterogeneity CLPs. L. Size distribution of CDCNs derived from β-sheet heterogeneity CLPs. Nanoparticle size slightly decreased with increased complexity of the N-terminal β-sheet domain. M. Drug release profiles for CDCNs derived from β-sheet heterogeneity CLPs. The release rate showed proportional correlation with the complexity of the N-terminal β-sheet domain. N. Encapsulation efficiency of CDCNs. The increase in the N-terminal β-sheet domain markedly reduced encapsulation efficiency.

TEM analysis revealed that CDCNs with larger α-helix domains (V_α666_CLP3-DOX) had thicker protein shells (∼5 nm) compared to those with smaller domains (V_α102_CLP3-DOX, ∼2 nm) (Fig. 3B). These findings suggest that the size of the rigid α-helix structure has a direct, positive correlation with the thickness of the nanoparticle shell without changing in zeta potential of the CDCNs (Fig. 3G). The larger α-helix domains likely provide a more extended and rigid structural framework, which results in a thicker protein shell around the nanoparticle core.

DLS analysis confirmed that nanoparticle size increased with increasing α-helix domain length (Fig. 3E). Although no significant differences in encapsulation efficiency were observed (Fig. 3C), the increasing of the α-helix domain notably reduced the release of DOX (Fig. 3F). The 50% release rate for V_α102_CLP3-DOX occurred within approximately 15 hours in pH=5 PBS buffer, while the release of V_α666_CLP3-DOX took longer than 40 hours— 4-times longer than V_α102_CLP3-DOX (Fig. 3F). This prolonged release likely results from the thicker protein shell in V_α666_CLP3-DOX, which reduces initial porosity and slows drug diffusion.

In contrast, we explored β-sheet domains by designing constructs of varying lengths, therefore three CLPs with different β-sheet N-terminal rigid domains were designed, denoted as V_β102_CLP3, V_β222_CLP3 and V_β666_CLP3, respectively (Fig. 3H, J, Fig. S2B). TEM images revealed that alterations in the β-sheet structure had minimal impact on the nanoparticle shell and zeta potential of CDCNs (Fig. 3I, K). Interestingly, increasing the length of the β-sheet domain slightly reduced nanoparticle size (Fig. 3L) and longer β-sheet domains decreased encapsulation efficiency (Fig. 3N). These findings may be attributed to the increased hydrophobicity of the protein carriers with the longer β-sheet domain, resulting in reduced binding of the hydrophobic drug.

Notably, and in contrast to the α-helix proteins, the increase in β-sheet structure significantly accelerated the release of DOX (Fig. 3M). This is likely due to the more rigid, planar conformation of β-sheets, which reduces internal resistance to drug diffusion, resulting in faster release compared to the flexible α-helical structures.

### DOX Conjugation Sites in CLPs Regulate the Properties of CDCNs

To further investigate the impact of chimeric structure variations in CLPs on the properties of CDCNs, we examined the effects of varying drug conjugation sites on CDCNs. Three CLPs with 1, 3, or 5 DOX conjugation sites were engineered, incorporating a rigid globular domain of 222 base pairs in an α-helix conformation and a flexible linear domain composed of 66 (Gly-Xaa-Yaa) repeats, based on prior findings (Fig. 4A, Fig. S3). TEM images showed that V_α222_CLP3GC5-DOX exhibited a significantly smaller size (∼150 nm) compared to V_α222_CLP3GC1-DOX (∼500 nm) (Fig. 4B). This finding was further supported by DLS data, which indicated that the size of the CDCNs decreased as the number of DOX conjugation sites increased (Fig. 4C). Specifically, for every additional two (GC) DOX conjugation sites, the particle size decreased by approximately 100 nm. Furthermore, the encapsulation efficiency increased by approximately 20% with the addition of two (GC) DOX conjugation sites (Fig. 4D). We hypothesize that the increase in DOX binding sites facilitates a higher binding capacity, leading to improved encapsulation efficiency. This, in turn, results in a denser CDCNs structure, which reduces the overall particle size. FTIR spectroscopy results indicate that the b-peak at 700 nm corresponds to the vibration of the S–C bond, representing the formation of thioether bonds (Fig. S4). A higher number of binding sites leads to a stronger b-peak intensity, indicating increased binding reaction efficiency. Drug release rates were slower in constructs with a higher number of conjugation sites, regardless of whether the pH was 5.0 or 7.4 (Fig. 4E). This phenomenon can be attributed to the increased drug loading within the nanoparticles as the number of conjugation sites rises. The aggregation of hydrophobic DOX molecules leads to the formation of a denser hydrophobic core, resulting in more homogeneous and stable CDCNs, thereby slowing the release rate of DOX.

**Figure 4.**
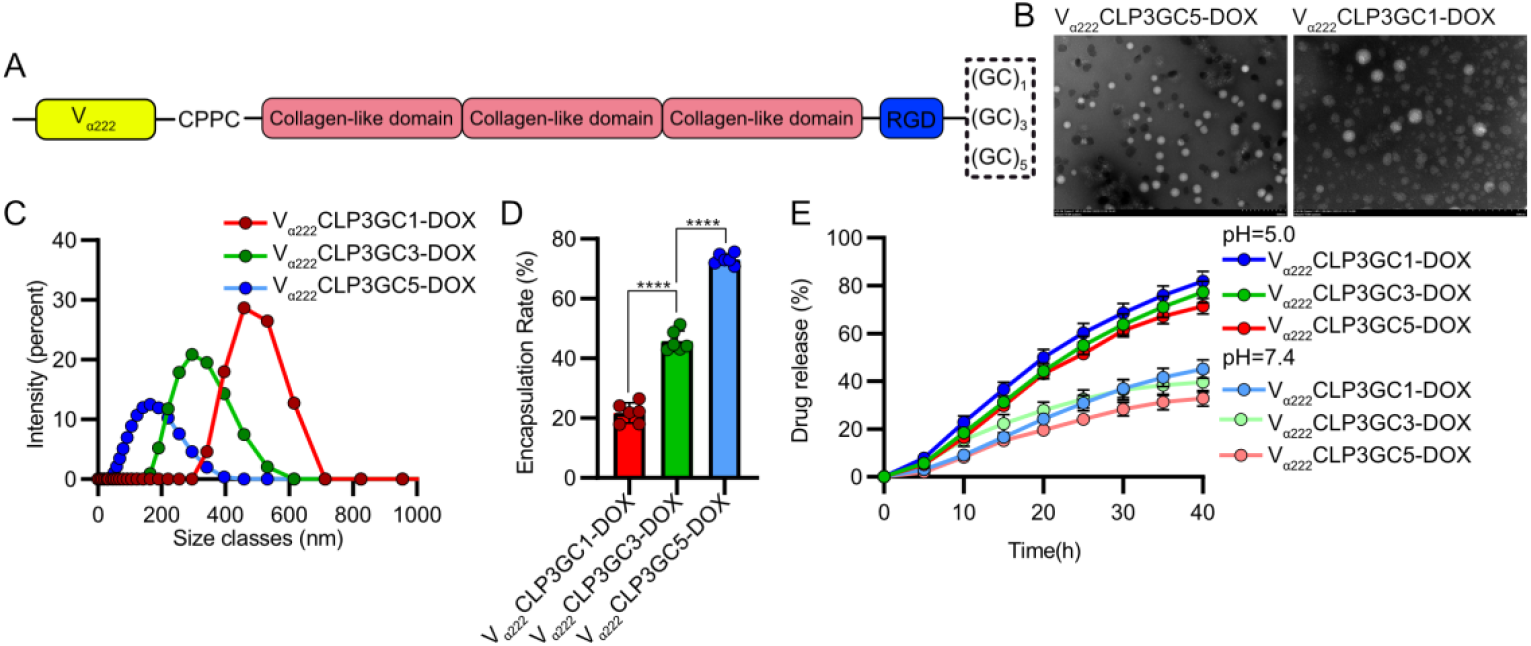
Impact of DOX Conjugation Sites on the Macrostructure and Drug Encapsulation of CDCNs. A. Schematic representation of the construction of CLPs with varying DOX conjugation sites. Activated DOX was conjugated with (GC) polypeptides, and 1, 3, or 5 DOX conjugation sites were genetically encoded at the C-terminal of the CLPs. B. TEM images of CDCNs prepared with V_α222_CLP3GC1-DOX and V_α222_CLP3GC5-DOX, showing the differences in nanoparticle size and morphology. C. DLS analysis of CDCNs with varying numbers of DOX conjugation sites, highlighting the size distribution of the nanoparticles. D. Encapsulation efficiency of CDCNs with different DOX conjugation site numbers. E. DOX release profiles of CDCNs in PBS at pH 5 and pH 7, demonstrating the effect of DOX conjugation site number on drug release kinetics.

### Anti-TNBC effects of N-terminal UPA modified CDCNs in vitro

We hypothesis that the V_α222_CLP3GC5 would be the most effective carrier for tumor targeting due to its optimal particle size, high DOX loading capacity, and moderate release rate. To verify this, we first co-cultured CDCNs prepared with different CLPs with 4T1 cells across a concentration gradient to determine the optimal drug concentration for subsequent experiments (Fig. S5). Subsequently, CDCNs prepared with different CLPs at a 25% concentration were co-cultured with HeLa cells to evaluate their antitumor efficacy. As expected, V_α222_CLP3GC5-DOX exhibited the most potent antitumor activity, inducing apoptosis in 63.9% of tumor cells after 12 hours of co-culture, a significantly higher rate than that observed with other formulations (Fig. 5B).

**Figure 5.**
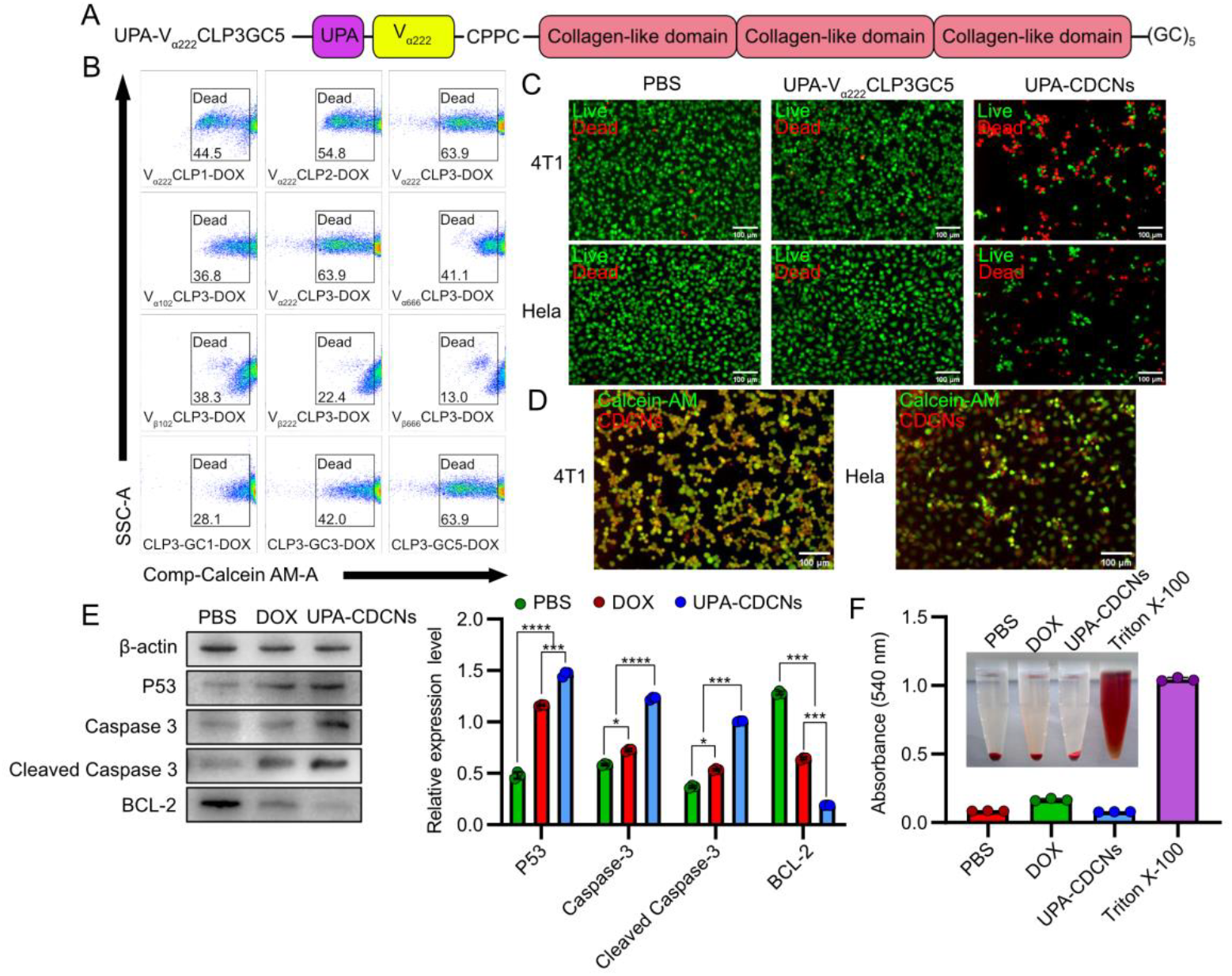
Anti-tumor Effects of TNBC-Targeting CDCNs in vivo. A. Schematic representation of the construction of TNBC-targeting CLPs. The TNBC-targeting peptide, UPA, was genetically encoded at the N-terminal of V_α222_CLP3GC5 to generate UPA-V_α222_CLP3GC5. B. Flow cytometry analysis of cell viability staining. The results demonstrate the anti-tumor efficacy of various CDCNs. V_α222_CLP3GC5-DOX showed the most pronounced tumor cell inhibition. C. Live/dead staining images to assess the anti-tumor effects of UPA-V_α222_CLP3GC5-DOX. D. Imaging of UPA-V_α222_CLP3GC5-DOX targeting ability. Cells were stained with Calcein AM and imaged in the FITC channel. Red fluorescence represents the spontaneous fluorescence of DOX, captured in the TRITC channel. E. Western blot analysis and quantitative statistics of 4T1 cells after treatment with PBS, DOX, and UPA-V_α222_CLP3GC5-DOX. F. Hemolysis assay images and quantitative data confirming the biocompatibility of UPA-V_α222_CLP3GC5-DOX.

We then hypothesized that modifying the N-terminal rigid structure with targeting peptides could further enhance the targeting and anti-tumor efficacy of the CDCNs. To test this hypothesis, we genetically encoded the TNBC-targeting peptide, UPA, to the N-terminal of V_α222_CLP3GC5, resulting in the creation of UPA-V_α222_CLP3GC5 (Fig. 5A, Fig. S6). UPA-V_α222_CLP3GC5-DOX was prepared and its targeting capability evaluated by co-culturing 5% UPA-CDCNs with TNBC 4T1 cells and HeLa cells. The results showed that UPA-CDCNs specifically co-localized with 4T1 cells, whereas in HeLa cells, UPA-CDCNs predominantly localized to the extracellular matrix (Fig. 5D), confirming its selective targeting of TNBC cells. Next, we increased the concentration of UPA-CDCNs to 25% and co-cultured with tumor cells for 24 hours to assess anti-tumor effects. UPA-CDCNs exhibited potent anti-tumor activity, inducing significant apoptosis (Fig. 5C). We further evaluated the expression of apoptosis-related proteins, including P53, Caspase-3, Cleaved Caspase-3, and BCL-2, as DOX induces apoptosis via the P53-dependent Caspase-3 pathway^31^. UPA-CDCNs treatment significantly upregulated P53, Caspase-3, and Cleaved Caspase-3 expression, while decreasing BCL-2 levels, compared to PBS and DOX-free controls (Fig. 5E), suggesting that UPA-CDCNs enhance DOX-induced apoptosis without altering its underlying mechanism. Finally, to assess the safety of UPA-CDCNs, a hemolysis test was performed. The results showed no hemolysis, confirming the biocompatibility of UPA-CDCNs (Fig. 5F).

### In Vivo Evaluation of UPA-CDCNs in TNBC Treatment

We evaluated the in vivo performance of UPA-CDCNs using an in-situ TNBC mouse model. After injecting 4T1-Luc cells into the mammary glands of mice and allowing tumor growth for 7 days, treatment was initiated once tumors reached 5 mm (Fig. 6A). To assess pharmacokinetics and targeting, Cy-7-labeled UPA-CDCNs and CDCNs were intravenously administered (Fig. 6D). Fluorescence imaging at 0, 8, 16, and 24 hours revealed prolonged tumor retention of ^Cy-7^UPA-CDCNs, with fluorescence persisting for up to 24 hours, compared to the rapid clearance of ^Cy-7^CDCNs by 16 hours (Fig. 6D). Regarding the major organs, ^Cy- 7^CDCNs was predominantly enriched in the heart and kidneys, whereas ^Cy-7^UPA-CDCNs showed robust fluorescence specifically at the tumor site (Fig. 6G). These findings suggest that the UPA-CDCNs prolongs the enrichment of CDCNs at the tumor site, enhancing tumor targeting.

**Figure 6.**
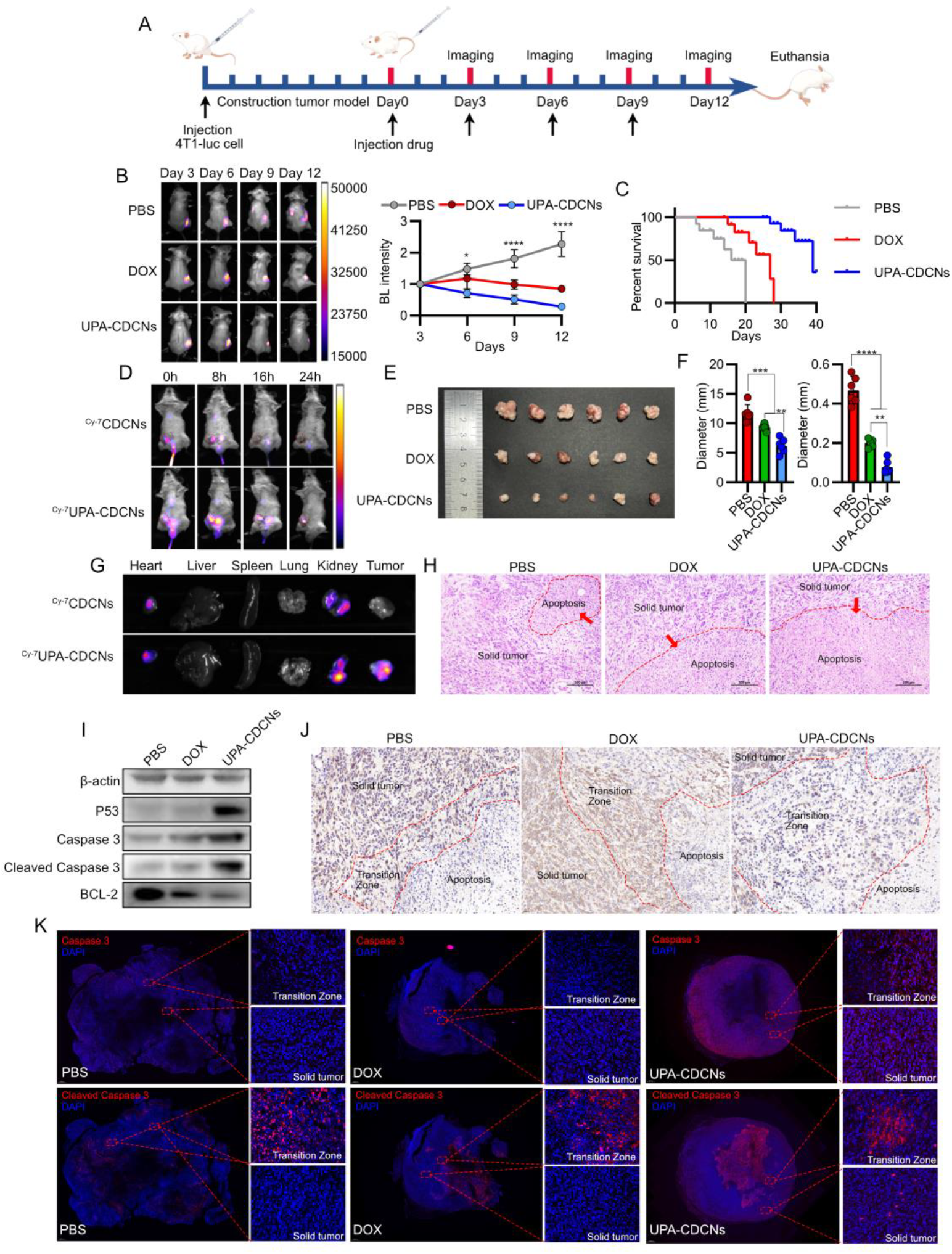
Anti TNBC Effects of UPS-CDCNs in Vivo. A. Experimental timeline for in vivo studies. The 4T1-Luc stable cell line was subcutaneously injected into mice, followed by a 7-day tumorigenic stage. Mice were then treated on Days 0, 3, 6, and 9, and imaged on Days 3, 6, 9, and 12. B. Bioluminescence imaging and quantification of relative bioluminescence intensity to monitor tumor progression in the TNBC mouse model. C. Survival curves of TNBC-bearing mice following different treatments. D. Fluorescence imaging of live mice to assess the pharmacokinetics of CDCNs and UPA-CDCNs. E. Representative tumor images following dissection of TNBC-bearing mice. F. Recorded tumor weight and diameter of solid tumors after treatment. G. Fluorescence imaging of major organs to investigate the biodistribution of CDCNs and UPA-CDCNs. H. H&E staining of tumor tissue sections, focusing on the central tumor region. Red arrows highlight areas of apoptosis in the solid tumor. I. Western blot analysis of protein expression in whole tumor tissues. J. Immunohistochemistry images showing BCL-2 expression in the central solid tumor. Red dashed areas indicate the apoptosis zone, solid tumor, and transition zone. K. Immunofluorescence images showing expression of Caspase 3 and Cleaved Caspase 3 throughout the tumor tissue.

TNBC modeled mice were treated with PBS, DOX, or UPA-CDCNs via intravenous injection on days 0, 3, 6, and 9 (Fig. 6A). Bioluminescence imaging showed a 50% reduction in tumor cell signals after UPA-CDCNs treatment, compared to only a 10% reduction with DOX and a 120% increase with PBS (Fig. 6B). The tumor size in the UPA-CDCNs-treated group remained between 3-6 mm, while it ranged from 11-14 mm in the PBS group and 8-10 mm in the DOX group (Fig. 6E, F). Tumor weight in the UPA-CDCNs group was 4.6-fold lower than PBS and 2.1-fold lower than DOX (Fig. 6E, F). UPA-CDCNs treatment also significantly extended survival, with a median of over 40 days, compared to 18 days for PBS and 28 days for DOX (Fig. 6C). These results suggest that UPA-CDCNs demonstrates superior efficacy as a DOX delivery system in the treatment of TNBC.

Histological analysis revealed significant apoptosis in the central regions of tumors treated with UPA-CDCNs, surpassing PBS and DOX groups (Fig. 6H). To further investigate apoptosis-related protein expression in the entire solid tumor, western blotting showed elevated expression of apoptosis markers, including Caspase-3, P53, Cleaved Caspase-3, and reduced BCL-2 in UPA-CDCNs-treated tumors (Fig. 6I, Fig. S8). To gain further insights into apoptosis protein expression, immunohistochemical and immunofluorescence staining was used to examine BCL-2, Caspase-3, and Cleaved Caspase-3 levels. Immunohistochemistry confirmed reduced BCL-2 in the transition zone (Fig. 6J), while immunofluorescence demonstrated increased expression of Cleaved Caspase-3, and Caspase-3 in both the transition zone and solid tumor (Fig. 6K). Collectively, these data suggest that UPA-CDCNs treatment promotes tumor apoptosis in both the transition zone and solid tumor. This effect may be attributed to the enhanced penetration ability of CDCNs facilitated by UPA modification.

### Reduced Biotoxicity of UPA-CDCNs Compared to Free DOX

Following the assessment of anti-TNBC efficacy, we next investigated the biocompatibility of CDCNs. Mice with breast cancer treated with PBS and free DOX showed notable spleen enlargement after the end of the experimental cycle of treatment, whereas no significant changes were observed in the spleen following treatment with UPA-CDCNs (Fig. 7A). To evaluate potential organ toxicity, we performed histological analysis of major organs, including the heart, liver, spleen, and kidneys. The results revealed that free DOX treatment induced cardiomyopathy, consistent with previous reports. However, no pathological changes were observed in the heart, liver, spleen, or kidneys of mice treated with UPA-CDCNs or PBS (Fig. 7E). These findings suggest that while DOX treatment induces myocardial injury, UPA-CDCNs do not cause similar cardiotoxicity or organ damage. Blood biomarkers were analyzed to further evaluate the safety of the treatments. UPA-CDCN treatment led to a slight reduction in alkaline phosphatase (ALP), a marker associated with splenomegaly. Additionally, UPA-CDCN-treated mice exhibited decreased levels of alanine aminotransferase (ALT, normal value: 13.5±5.3 U/L) and aspartate aminotransferase (AST, normal value: 36.2±6.13 U/L), bringing them closer to normal values and suggesting a reduction in liver function abnormalities. In contrast, DOX treatment resulted in a significant increase in serum creatine kinase (CK) and lactate dehydrogenase (LDH-L) levels, both of which are markers of myocardial damage.

**Figure 7.**
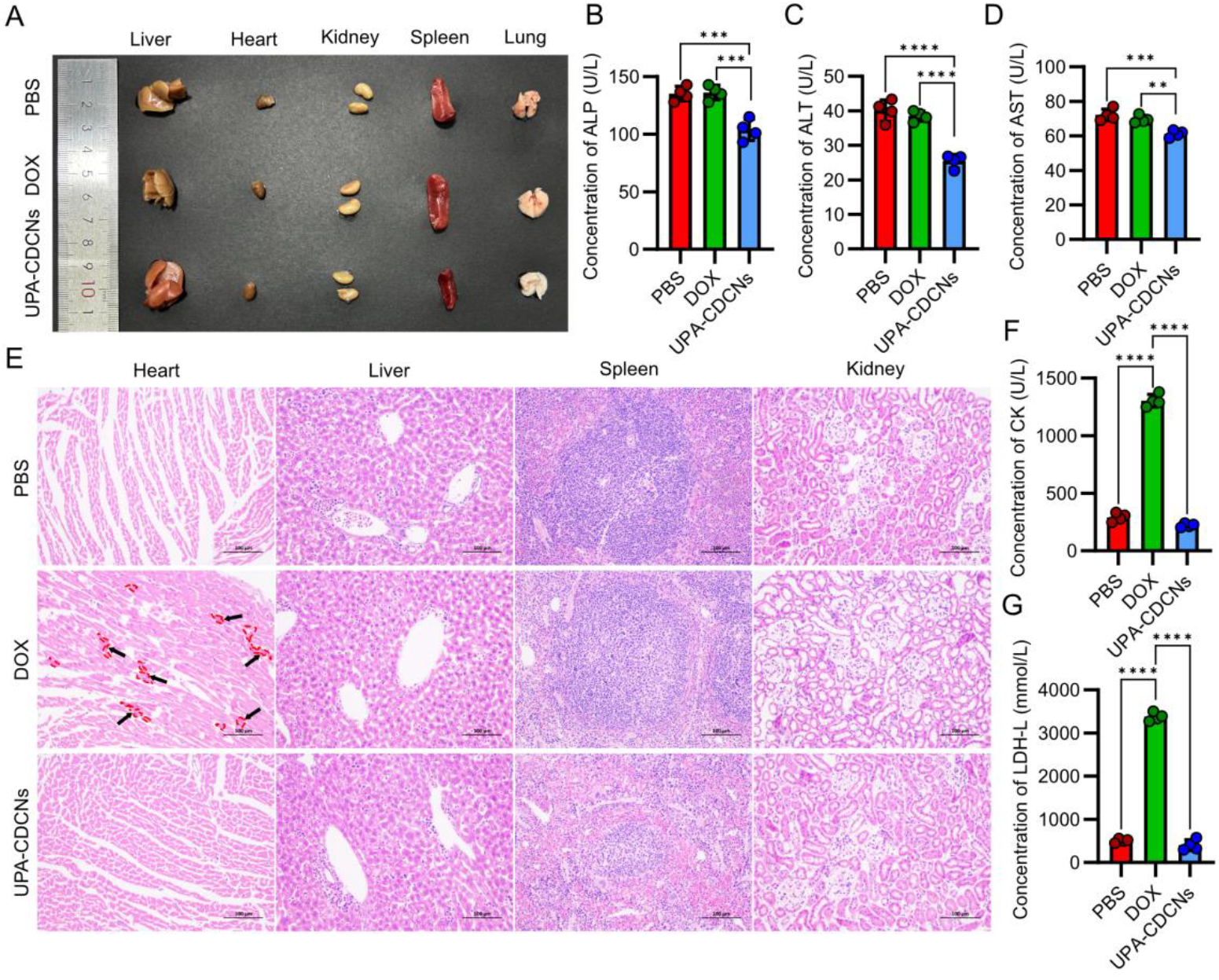
Reduced Biotoxicity of UPA-CDCNs Compared to Free DOX. A. Photographs were taken of various tissues of TNBC mice treated with PBS, DOX and UPA-CDCNs. B. Measurement of alkaline phosphatase (ALP) in serum from TNBC mice. Enlarged spleen resulted in elevated alkaline phosphatase. C. Measurement of alanine transaminase (ALT) in serum from TNBC mice. D. Measurement of aspartate aminotransferase (AST) in serum from TNBC mice. E. H&E staining of heart, liver, spleen and kidney tissues from TNBC mice. Red dashed lines and black arrows emphasize free DOX-induced cardiomyopathy shown as dark punctate masses in cardiac sections. F. Measurement of creatine kinase (CK) in serum from TNBC mice. G. Measurement of lactate dehydrogenase (LDH-L) in serum from TNBC mice.

### LinTT1 Modified CDCNs cross blood-brain barrier (BBB) and Anti-GBM Effects

The above results demonstrate that the V_α222_CLP3GC5-based CDCNs exhibit effective targeting and anti-TNBC activity with UPA modification. We further investigated whether V_α222_CLP3GC5 could serve as a brain delivery system for GBM therapy. To test this hypothesis, we genetically engineered a P32 targeting peptide, LinTT1, as a GBM targeting ligand^28^, at the N-terminal of V_α222_CLP3GC5, resulting in the construction of LinTT1-V_α222_CLP3GC5 (Fig. S9, Fig. S10A). After expression, purification, and conjugation with DOX, LinTT1-CDCNs were prepared and their anti-GBM effects were evaluated in vitro and in vivo.

We first assessed the anti-GBM efficacy of LinTT1-CDCNs using the GL261 cell line. Treatment with 25% LinTT1-CDCNs for 12 hours resulted in a 64.3% cell death rate, significantly higher than the 19.8% observed in the DOX-only group (Fig. S10B). To investigate the mechanism of action, we analyzed the P53-dependent Caspase-3-mediated apoptotic pathway^31^. Western blot analysis revealed that LinTT1-CDCNs treatment significantly upregulated the expression of P53, Caspase-3, Cleaved Caspase-3, and tumor necrosis factor-α (TNF-α), while downregulating BCL-2 expression (Fig. S10E). These results suggest that LinTT1-CDCNs promotes GBM cell apoptosis through activation of the P53-Caspase cascade and modulation of key apoptotic regulators.

We further investigated the anti-GBM effects of LinTT1-CDCNs in vivo, with an emphasis on its ability to cross the BBB. In vitro transwell assays showed that LinTT1-CDCNs crossed the BBB model, as indicated by robust red fluorescence in the LinTT1-CDCNs group, whereas the DOX-only group showed no fluorescence (Fig. S11A). For the in vivo investigation, we first established a GBM mouse model via brain stereotactic injection of the GL261-Luc stable cell line^32^ (Fig. S11B). LinTT1-CDCNs and CDCNs were both conjugated with Cy-7 and intravenously injected into mice (Fig. 8B). In vivo imaging of Cy-7-labeled LinTT1-CDCNs (^Cy-7^LinTT1-CDCNs) and CDCNs (^Cy-7^CDCNs) revealed that LinTT1-CDCNs successfully crossed the BBB, with fluorescence detected in the brain of treated mice at 24 hours post-injection (Fig. 8C). In contrast, fluorescence in the ^Cy-7^CDCNs group was limited to the heart and kidneys (Fig. 8C). These findings confirm that LinTT1-modified CDCNs can efficiently target GBM and penetrate the BBB.

**Figure 8.**
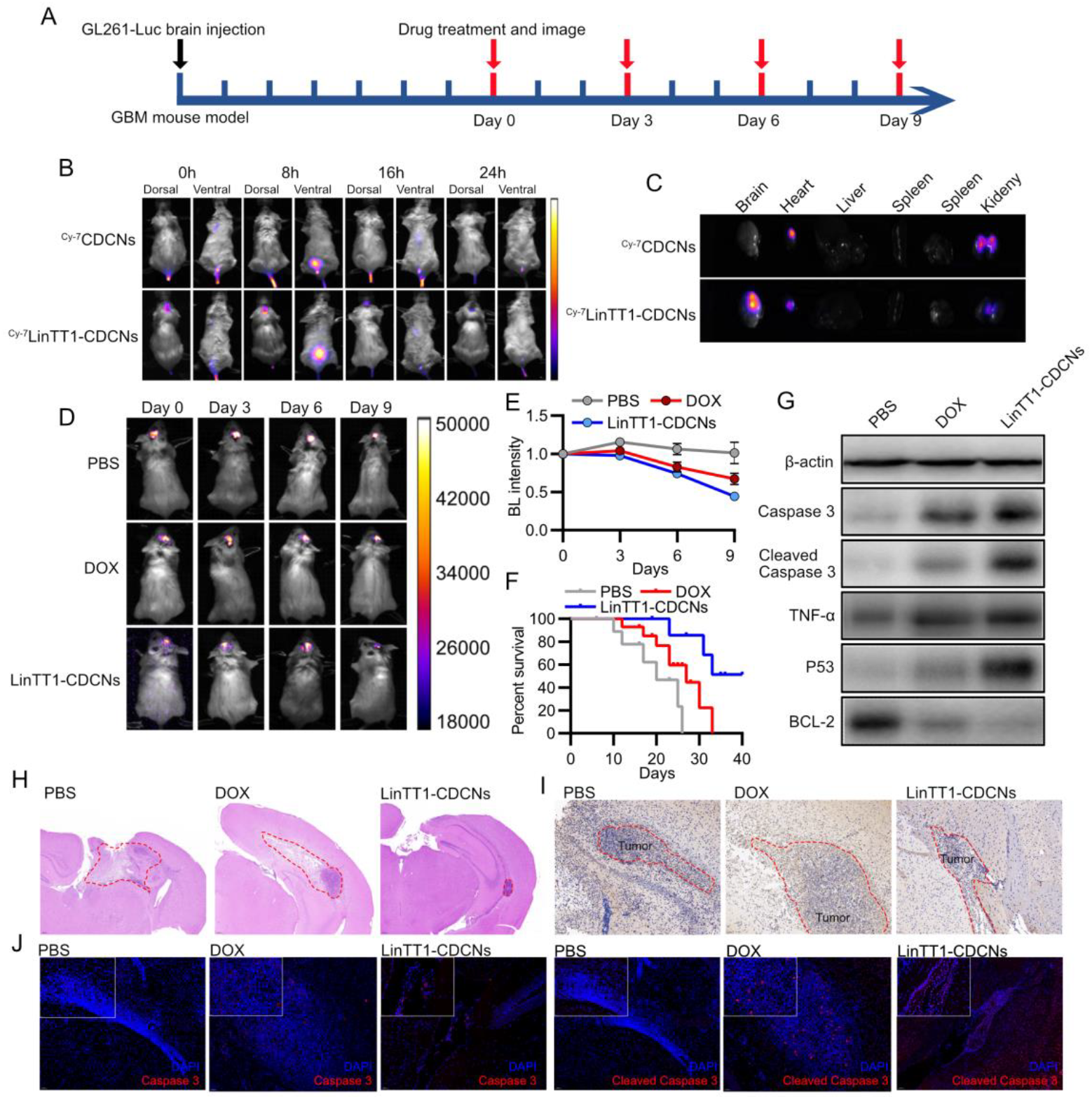
Anti-GBM Effects of LinTT1 modified CDCNs. A. Experimental timeline for evaluating the anti-GBM effects of LinTT1-CDCNs. The GL261-Luc stable cell line was used to establish the GBM mouse model, followed by a 7-day tumorigenic stage. Mice were then administered treatments and imaged on days 0, 3, and 6. B. Fluorescence imaging of live mice to assess the pharmacokinetics of CDCNs and LinTT1-CDCNs. Both were labeled with Cy-7 fluorescence dye for tracking. C. Fluorescence imaging of major organs to assess the biodistribution of CDCNs and LinTT1-CDCNs 24 hours post-treatment. D. Bioluminescence imaging of GBM model mice following treatment with LinTT1-CDCNs, DOX, and PBS. E. Relative bioluminescence intensity quantification of GBM following different treatments. F. Survival curves of GBM model mice following different treatments. G. Western blot analysis of apoptosis-related protein expression in the whole brain of GBM model mice after treatment. H. H&E staining of brain tissue from the GBM mouse model. The red dashed area indicates the GBM site. I. Immunohistochemistry images showing BCL-2 expression at the GBM site. J. Immunofluorescence staining showing the expression of Caspase-3 and Cleaved Caspase-3 at the GBM site following treatment.

For in vivo anti-GBM efficacy, GL261-Luc cells were stereotactically injected into the brains of mice. LinTT1-CDCNs were intravenously administered on Days 0, 3, 6, and 9 (Fig. 8A). Bioluminescence imaging showed a 50% reduction in tumor bioluminescence in the LinTT1-CDCNs-treated group, compared to a 20% reduction in the DOX group and a 20% increase in the PBS group (Fig. 8E). Survival was also significantly prolonged in the LinTT1-CDCNs group, with a median survival of over 40 days, compared to 18 days for PBS and 26 days for DOX (Fig. 8F).

After 12 days, brain tissue was analyzed for pathological changes. H&E staining of brain tissue revealed that treatment with LinTT1-CDCNs significantly reduced the size of the GBM site compared to the DOX and PBS treatment groups (Fig. 8H). Western blot analysis confirmed increased expression of P53, Caspase-3, Cleaved Caspase-3, and TNF-α in the LinTT1-CDCNs group (Fig. 8G, Fig. S13). Immunohistochemistry and immunofluorescence revealed that LinTT1-CDCNs treatment decreased BCL-2 expression and significantly increased Cleaved Caspase-3 expression at the tumor site compared to both DOX and PBS treatments (Fig. 8I, J). These results suggest that LinTT1-CDCNs promote apoptosis in GBM tumors more effectively than free DOX.

## Discussion

In this study, we introduce a novel chimeric CLP-like protein design strategy, constructing a series of structurally heterogeneous chimeric proteins. By leveraging these engineered proteins, we successfully conjugated doxorubicin (DOX) to create CDCNs. The inherent tunability of the protein structure facilitated precise control over the macrostructure and functional properties of the CDCNs. Furthermore, by genetically encoding a targeting peptide at the N-terminal of the designed proteins, we achieved targeted delivery of the CDCNs to both TNBC and GBM. The CDCNs not only enhanced the anti-tumor efficacy of DOX but also significantly mitigated the typical side effects associated with its use. Collectively, these findings highlight the potential of chimeric CLPs as an advanced protein-based drug delivery system, with the added advantage of precise control over delivery properties through structural heterogeneity.

Protein-based drug delivery systems have emerged as a promising alternative to chemical polymers due to their natural biocompatibility and versatility^7, 17^. However, the immunogenicity of proteins remains a significant concern regarding their safety and efficacy as drug delivery vehicles^33^. Natural proteins, such as bovine serum albumin (BSA), human serum albumin (HSA), and elastin, carry inherent risks due to their complex functional groups, which may trigger immune responses when used in drug delivery applications^1, 4, 33^. In contrast, genetically engineered de novo-designed CLPs minimize these risks by reducing the number of exposed functional groups, thus functioning as “blank-template proteins”^22, 26^. The production of these proteins through genetic engineering techniques ensures high purity, modularity, batch-to-batch reproducibility, and homogeneity, making them superior drug delivery carriers compared to natural proteins^1^.

One of the key challenges in protein-based drug delivery is relating the macroscopic properties of the material to its underlying microstructure. The inherent complexity of tertiary protein structures often leads to issues such as poor homogeneity and random aggregation, which can result in undesired macrostructures^1, 3^. Globular structural proteins, commonly used as drug carriers, face this issue, leading to instability and inefficient drug release profiles. In contrast, linear structural proteins, such as elastin and collagen, form less predictable “woolen-ball”-like structures due to hydrophobic molecular interactions, limiting their ability to provide insight into the relationship between protein structure and macroscopic properties^14, 20^. Chimeric proteins that combine both globular structural terminal domains and flexible linear sequences may offer a solution to this challenge, providing more uniform and stable protein-based carriers with improved control over both structure and function^14, 20^.

Our findings demonstrate that an increase in α-helix content resulted in a thicker CDCN shell, while the incorporation of linear structures led to a reduction in particle size. We hypothesize that the relationship between protein structure and macrostructure of the carriers is governed by two key mechanisms: the “barrier effect” and the “hydration-driven densification effect.” The “barrier effect” refers to the rigidity of the α-helix structure, which prevents folding and promotes the assembly of rigid α-helices, leading to the formation of a stable “shell” structure. In contrast, the “hydration-driven densification effect” arises from the introduction of hydrophilic groups in linear structures, which enhances intermolecular interactions within the nanoparticle. These interactions help reduce internal porosity, resulting in a more compact and stable nanoparticle structure with reduced particle size. These theoretical insights provide a foundation for further exploring how protein structure influences the macroscopic properties of protein-based materials.

Although a series of structurally heterogeneous CLPs have been successfully used to construct protein-based nanoparticles, their stimulus-responsive capabilities remain limited. To further enhance their functionality, stimulus-responsive domains, such as metalloproteinase cleavage sites^34^ or photodynamic response protein^35^, can be genetically integrated into the N-terminal of linear collagen-like sequences. This strategy would enable the development of dynamic, stimulus-responsive protein-based drug delivery systems. Moreover, the inherent functional diversity of natural proteins offers the potential to easily create multi-stimulus responsive carriers through genetic fusion, allowing for the combination of targeted drug delivery and stimulus-triggered release mechanisms^4, 5, 10^. These advances could significantly enhance the anti-tumor efficacy of protein-based nanoparticles and open new avenues for cancer therapy and other medical applications.

The targeting ability of CDCNs was engineered by genetically encoding a ligand peptide at the N-terminal of the rigid globular domain^28, 36^. Native CDCNs, in their unmodified form, lacked inherent targeting specificity, as demonstrated by our results. The incorporation of N-terminal modifications significantly enhanced the targeting efficiency, tumor permeability, and biodistribution of the CDCNs, highlighting the importance of genetic modifications for optimizing drug delivery properties. Furthermore, the genetic fusion of additional biofunctional peptides, such as immune-modulating peptides^37^ or photodynamic/photothermal peptides^35, 38^, provides a versatile platform for enhancing therapeutic outcomes. This modular approach offers a promising strategy for the development of multifunctional protein-based drug delivery systems with improved anti-tumor activity and additional therapeutic functions.

The structural heterogeneity of the designed CLPs allowed for the precise control of CDCN properties, including their macrostructure and drug release kinetics. The release profile was tunable, ranging from 20 to 40 hours, while particle size could be adjusted from 80 nm to 800 nm. These findings demonstrate the potential for designing protein carriers tailored to the specific requirements of different tumor treatments. Moreover, combining CDCNs with distinct properties could further enhance their therapeutic potential, not only for anti-tumor applications but also for other medical purposes, such as immune adjuvants^39^ or antibiotic delivery. The versatility of CLP-based carriers, achieved through the design and modification of chimeric proteins, offers immense potential for expanding their applications. In this study, we primarily focused on demonstrating the utility of structural heterogeneity in CLPs for targeted DOX delivery, but future research may explore the broader applications of these carriers in various therapeutic contexts.

## Methods

### Materials and reagents

Restriction endonucleases, DNA polymerase, and T4 DNA ligase were purchased from Thermo Scientific (Waltham, MA, USA). The hematoxylin-eosin (HE) staining kit was obtained from Solarbio (Shanghai, China). Phosphate-buffered saline (PBS), DMEM, RPMI 1640, DMEM/F-12 media, trypsin, D-luciferin potassium, penicillin, and streptomycin were purchased from Meilunbio (China). Fetal bovine serum (FBS) was obtained from DCELL. Doxorubicin (DOX) was purchased from MACKLIN Biochemical Technology Co. BMPH was supplied by Bidet Pharmaceuticals. Cy-7 fluorescent labeling protein was acquired from Xi’and Ruixi Biotech. The Calcein-AM/PI double staining kit was purchased from Beijing Solarbio Biotechnology Co. Matrix gel was obtained from Mogengel Bio. Skimmed milk was sourced from BD (USA). Primary antibodies were obtained from Abcam (USA), including anti-His (CAS: Ab18184), β-actin (CAS: Ab8226), P53 (CAS: Ab131442), Caspase-3 (CAS: Ab32499), Cleaved Caspase-3 (CAS: Ab32042), TNF-α (CAS: Ab183218), and CD63 (CAS: Ab216130). The BCL-2 antibody (CAS: GB124830) was purchased from Service Bio Co. Primers were synthesized by Sangon Biotech (Zhengzhou, China). BALB/c mice (6–8 weeks old, 24–26 g) were obtained from SPF (Beijing) Biotechnology Co.

### Plasmid Construction

Recombinant proteins were generated from lab-engineered plasmids containing collagen-like protein sequences derived from *Streptococcus pyogenes* (Scl-2). The structural rigid domain and linear collagen-like domain were designed in-house. To enhance targeting and membrane penetration, the UPA(VSNKYFSNIHWGC)^36^/ LinTT1(AKRGARSTA)^28^ peptide was incorporated at the N-terminus of the recombinant protein. Additionally, an RGD cell adhesion peptide and (GC)_n_ unit were introduced at the C-terminus. The gene construct was inserted into the pET-28a(+) vector (Addgene plasmid #90473) using T4 DNA ligase. The recombinant plasmid was then transformed into *E. coli* DH5α competent cells, plated on kanamycin-selective solid medium, and single colonies were isolated. The plasmid was extracted from expanded cultures and validated through sequencing to confirm successful fusion plasmid construction^25, 40^.

### Protein expression and purification

The recombinant plasmid was first transformed into E. coli Rosetta cells. The transformed bacteria were inoculated into LB medium supplemented with kanamycin and chloramphenicol and incubated at 37°C with shaking. When the optical density at 600 nm (OD_600_) reached 0.6–0.7, isopropyl β-D-1-thiogalactopyranoside (IPTG, 0.3 mM) was added to induce protein expression. The culture was maintained at 16°C for 24 h before harvesting the bacterial cells by centrifugation. The bacterial pellet was lysed, and the supernatant was collected, resuspended, and processed to minimize aggregation. The lysate was then subjected to Ni^2+^-NTA affinity chromatography for protein purification. The purified protein was analyzed for size and purity using SDS-PAGE followed by Coomassie blue staining, with further validation by Western blotting using an anti-His antibody. The eluted protein was concentrated to 2.5 mg/mL using a 10-kDa molecular weight cutoff centrifugal filter at 3,000 rpm and stored for further use^24, 40^.

### Preparation of CDCNs

BMPH (0.025 g) and DOX (0.055 g) were each dissolved in 5 mL of methanol. The BMPH solution was then added dropwise into the DOX solution under continuous stirring, and the reaction was allowed to proceed for 12 h to form BMPH-DOX. To synthesize CDCNs, 110 μL of BMPH-DOX and 1 mL of purified recombinant protein (2.5 mg/mL) were mixed in 10 mL of PBS (pH 7.4). The mixture was stirred for 12 h to obtain CDCNs with a final BMPH-DOX concentration of 1%. Free DOX was prepared separately by dissolving 0.001 g of DOX in 10 mL of water and stored at 4°C^14, 15, 20^.

### Characterization of CDCNs

To evaluate the properties of CDCNs, cumulative DOX release was assessed by placing 5 mL of CDCNs in a dialysis bag submerged in 50 mL PBS (pH 5.0, pH 7.4), with absorbance at 480 nm measured at 1-hour intervals using UV spectrophotometry. The cumulative release rate (CRr) was calculated as CRr = (∑4Y_t_/TD) × 100%. Encapsulation efficiency was determined by mixing 10 μL of BMPH-DOX with PBS (pH 7.4), measuring absorbance at 480 nm, and calculating efficiency as [(Ct - C_0_)/Ct] × 100%. For morphological analysis, CDCNs were dispersed in PBS, drop-cast onto a copper mesh, dried, and imaged using an HT 7800 transmission electron microscope at 120 kV. Particle size, polydispersity index (PDI), and zeta potential were measured via dynamic light scattering (DLS) after diluting CDCNs 5-fold in ultrapure water. Fourier Transform Infrared Spectroscopy (FTIR) was performed to obtain spectral profiles (400–4000 cm^−1^) using the diffuse reflectance method.

### Cellular experiment

4T1, GL261, NIH-3T3, HeLa, and U87MG cells were cultured in appropriate media supplemented with 10% FBS and 1% penicillin-streptomycin at 37°C in a 5% CO2 incubator. To assess cell viability, 4T1 cells were treated with varying concentrations of CDCNs (5%, 15%, 25%, 35%) for 12 hours and stained with Calcein-AM and PI. The number of dead cells increased with higher CDCN concentrations. Based on these results, 25% UPA-CDCNs were used for live/dead staining of 4T1 and HeLa cells, and 25% LinTT1-CDCNs for U87MG and HeLa cells. Drug targeting was assessed by treating cells with UPA-CDCNs (4T1 and HeLa) or LinTT1-CDCNs (U87MG and HeLa), followed by Calcein-AM staining and fluorescence microscopy. For the transwell assay, NIH-3T3 cells were cultured in the upper chamber and GL261 cells in the lower chamber. After 12 hours, cells were stained with Calcein-AM. For flow cytometry, 15% CDCNs were applied to HeLa cells for 12 hours, followed by Calcein-AM staining and analysis. The most lethal CDCNs were identified for HeLa and GBM cells.

### Animal experiments on GBM mice

Balb/c mice (6-8 weeks old, 24-26 g) were acclimatized for 7 days, and GL261-Luc cells were resuspended in PBS, mixed with matrix gel (1:1), and injected into the right intracranial coronal region of the striatum caudate nucleus using a stereotactic apparatus. After anesthesia with isoflurane, a 1-mm diameter hole was drilled in the skull, and 5μL of the cell suspension was injected at a rate of 5μL/min to deliver 2.5×10^5^ cells. After the procedure, the mice were monitored for recovery. Six days later, D-luciferin potassium was injected for bioluminescence imaging. For the GBM animal study, after model establishment (day 0), mice were divided into groups and administered drugs every 3 days. Imaging was conducted on the same schedule for a total of 5 sessions. Mice were euthanized on day 21, and brains and tissues were dissected for protein extraction and fixation (4% PFA). **In vivo targeting assay of Cy-7 fluorescently labeled CDCNs**. After successfully constructing mouse models of breast cancer and glioma, 1 ml of each of the three protein nanocomplexes including CDCNs, UPA-CDCNs, and LinTT1-CDCNs, was added with 100μL of coupling reagent and 0.5 mg of Cy7-mixture (Excitation 745nm, Emission 800nm), mixed well, and incubated for 30 min in the dark. The reaction solution was then transferred to an ultrafiltration tube, centrifuged at 10,000 rpm for 5 min to remove the free dye, and made up to 1 ml with PBS to resuspend the truncated drug. The two animal models were divided into control and experimental groups, the control group was injected with CDCNs, UPA-targeted mice with breast cancer, and LinTT1-targeted mice with glioma. Each mouse was injected with 100μL in the tail vein, and the injection was recorded as 0 h. The fluorescence distribution was observed by using 740 nm excitation light, and imaged at the 0h, 8h, 16h, and 24h. After the imaging was completed, the mouse was dissected, and its tissues were dissected out and fluorescence imaging was performed to observe the drug targeting.

### Histological analysis

Mice underwent cardiac perfusion, and tissues (brain, heart, liver, spleen, kidney, and tumors) were collected and fixed in 4% PFA. After 24 hours, tissues were sequentially dehydrated in graded ethanol (30%, 50%, 70%, 80%, 95%, and 100%), cleared in xylene, and embedded in paraffin. Paraffin blocks were sectioned at 4 μm, spread, and baked. Sections were deparaffinized in xylene, rehydrated through graded ethanol, stained with hematoxylin, differentiated, counterstained with eosin, dehydrated, permeabilized, mounted, and imaged.

### Hemolysis experiment

Fifteen percent whole blood from TNBC mice was incubated with PBS, DOX, UPA-CDCNs, or Triton X-100 at 37°C for 1 hour. Samples were then centrifuged at 1,500 rpm for 10 minutes, photographed, and 200 μL of the supernatant was collected for OD540 measurement.

### Immunohistochemical and Immunofluorescence Analysis

4T1 and GL261 cells were treated with PBS, DOX, or CDCNs for 24 h, lysed in buffer with phosphatase and protease inhibitors, and centrifuged at 12,000 rpm for 30 min at 4°C. The supernatant was collected for Western blot analysis using β-actin as a loading control. Glioma-bearing brains and breast tumors were similarly processed for Western blotting with β-actin, P53, Caspase-3, Cleaved Caspase-3, and BCL-2. For tissue immunofluorescence, paraffin sections were deparaffinized, rehydrated, subjected to antigen retrieval, and blocked with 1% BSA. Primary antibodies (Caspase-3, Cleaved Caspase-3) were incubated overnight at 4°C, followed by secondary antibody incubation, DAPI and SpOrange staining, and mounting. Immunohistochemistry followed a similar deparaffinization and permeabilization process, with endogenous peroxidase blocked using 3% H2O2. After primary (BCL-2) and secondary antibody incubation, streptavidin-HRP was applied, followed by DAB development, hematoxylin counterstaining, dehydration, and mounting.

### Statistical analysis

Data analysis was performed using Prism 10.1.2 and Origin 2021. Statistical significance was determined using two-way ANOVA for multiple group comparisons (*p < 0.05, **p < 0.01, ***p < 0.001, ****p < 0.0001), with results presented as the standard error of the mean (SEM). Image processing was conducted using Fiji software.

## Acknowledgement

This project was supported by a grant from National Natural Science Foundation (22365022); Inner Mongolia Natural Science Foundation (2022QN03014); Inner Mongolia Science and Technology Project (2022ZY0050); Inner Mongolia Youth Science and Technology Talent Support Project (NUYT23092); Science and Technology Leading Talent Team in Inner Mongolia Autonomous Region (2022LJRC0009); Inner Mongolia Autonomous Region Science and Technology Project (2023YFHH0010). All authors have approved the final version of this manuscript. We wish to thank the Electron Microscopy Centre of Inner Mongolia University for the microscopy and microanalysis of our specimens. Ethical approval for the in vivo diabetic wound healing experiments was granted by the Institutional Animal Care and Use Committee of Inner Mongolia University.

## Conflict of Interest

The authors declare no conflict of interest.

## Data Availability Statement

The data supporting the findings and all plasmids and proteins are available upon reasonable request from the corresponding author.

## Supporting Information

The Supporting Information is available free of charge at: Supporting information.

Mass spectrum of peptide Q11, SDS-PAGE, Protein identity, CD spectroscopy, Live/dead staining images, Predict the 3D structure of proteins and Genetic sequence.

## Notes

### Competing Interest Statement

The authors have declared no competing interest.

## References

(1) Habibi, N.; Mauser, A.; Ko, Y.; Lahann, J. Protein Nanoparticles: Uniting the Power of Proteins with Engineering Design Approaches. Adv Sci (Weinh) 2022, 9 (8), e2104012. DOI: 10.1002/advs.202104012 From NLM Medline.

(2) Wu, S. Y.; Ye, Y. X.; Zhang, Q.; Kang, Q. J.; Xu, Z. M.; Ren, S. Z.; Lin, F.; Duan, Y. T.; Xu, H. J.; Hu, Z. Y.; et al. Multifunctional Protein Hybrid Nanoplatform for Synergetic Photodynamic-Chemotherapy of Malignant Carcinoma by Homologous Targeting Combined with Oxygen Transport. Adv Sci (Weinh) 2022, e2203742. DOI: 10.1002/advs.202203742 From NLM Publisher.

(3) Shi, M.; McHugh, K. J. Strategies for overcoming protein and peptide instability in biodegradable drug delivery systems. Advanced Drug Delivery Reviews 2023, 114904.

(4) Varanko, A.; Saha, S.; Chilkoti, A. Recent trends in protein and peptide-based biomaterials for advanced drug delivery. Advanced drug delivery reviews 2020, 156, 133–187.

(5) Wang, Z.; Zhang, S.; Zhang, R.; Chen, X.; Sun, G.; Zhou, M.; Han, Q.; Zhang, B.; Zhao, Y.; Jiang, B.; et al. Bioengineered Dual-Targeting Protein Nanocage for Stereoscopical Loading of Synergistic Hydrophilic/Hydrophobic Drugs to Enhance Anticancer Efficacy. Advanced Functional Materials 2021, 31 (29), 2102004. DOI: 10.1002/adfm.202102004.

(6) Mout, R.; Bretherton, R. C.; Decarreau, J.; Lee, S.; Gregorio, N.; Edman, N. I.; Ahlrichs, M.; Hsia, Y.; Sahtoe, D. D.; Ueda, G. De novo design of modular protein hydrogels with programmable intra- and extracellular viscoelasticity. Proceedings of the National Academy of Sciences 2024, 121 (6), e2309457121.

(7) Kianfar, E. Protein nanoparticles in drug delivery: animal protein, plant proteins and protein cages, albumin nanoparticles. Journal of Nanobiotechnology 2021, 19 (1), 159.

(8) Hassanzadeh, S.; Nematollahzadeh, A.; Mirzayi, B.; Fatemeh Kaboli, S. Protein-based nanoparticles synthesized at a high shear rate and optimized for drug delivery applications. Journal of Molecular Liquids 2021, 335. DOI: 10.1016/j.molliq.2021.116133.

(9) Liu, F.; Xue, L.; Xu, L.; Liu, J.; Xie, C.; Chen, C.; Liu, Y. Preparation and characterization of bovine serum albumin nanoparticles modified by Poly-l-lysine functionalized graphene oxide for BMP-2 delivery. Materials & Design 2022, 215. DOI: 10.1016/j.matdes.2022.110479.

(10) Bae, S.; Ma, K.; Kim, T. H.; Lee, E. S.; Oh, K. T.; Park, E. S.; Lee, K. C.; Youn, Y. S. Doxorubicin-loaded human serum albumin nanoparticles surface-modified with TNF-related apoptosis-inducing ligand and transferrin for targeting multiple tumor types. Biomaterials 2012, 33 (5), 1536–1546. DOI: 10.1016/j.biomaterials.2011.10.050 From NLM Medline.

(11) Gregory, J. V.; Kadiyala, P.; Doherty, R.; Cadena, M.; Habeel, S.; Ruoslahti, E.; Lowenstein, P. R.; Castro, M. G.; Lahann, J. Systemic brain tumor delivery of synthetic protein nanoparticles for glioblastoma therapy. Nat Commun 2020, 11 (1), 5687. DOI: 10.1038/s41467-020-19225-7 From NLM Medline.

(12) Mauser, A.; Waibel, I.; Banerjee, K.; Mujeeb, A. A.; Gan, J.; Lee, S.; Brown, W.; Lang, N.; Gregory, J.; Raymond, J.; et al. Controlled Delivery of Paclitaxel via Stable Synthetic Protein Nanoparticles. Advanced Therapeutics n/a (/a), 2400208. DOI: 10.1002/adtp.202400208.

(13) Song, Y.; Zhu, W.; Wang, Y.; Deng, L.; Ma, Y.; Dong, C.; Gonzalez, G. X.; Kim, J.; Wei, L.; Kang, S.-M.; et al. Layered protein nanoparticles containing influenza B HA stalk induced sustained cross-protection against viruses spanning both viral lineages. Biomaterials 2022, 287. DOI: 10.1016/j.biomaterials.2022.121664.

(14) Han, W.; Chilkoti, A.; Lopez, G. P. Self-assembled hybrid elastin-like polypeptide/silica nanoparticles enable triggered drug release. Nanoscale 2017, 9 (18), 6178–6186. DOI: 10.1039/c7nr00172j From NLM Medline.

(15) MacKay, J. A.; Chen, M.; McDaniel, J. R.; Liu, W.; Simnick, A. J.; Chilkoti, A. Self-assembling chimeric polypeptide-doxorubicin conjugate nanoparticles that abolish tumours after a single injection. Nat Mater 2009, 8 (12), 993–999. DOI: 10.1038/nmat2569 From NLM Medline.

(16) Bidwell, G. L., 3rd. Novel Protein Therapeutics Created Using the Elastin-Like Polypeptide Platform. Physiology (Bethesda) 2021, 36 (6), 367–381. DOI: 10.1152/physiol.00026.2021 From NLM Medline.

(17) Papi, M.; Palmieri, V.; Maulucci, G.; Arcovito, G.; Greco, E.; Quintiliani, G.; Fraziano, M.; De Spirito, M. Controlled self assembly of collagen nanoparticle. Journal of Nanoparticle Research 2011, 13 (11), 6141–6147. DOI: 10.1007/s11051-011-0327-x.

(18) Ragothaman, M.; Kannan Villalan, A.; Dhanasekaran, A.; Palanisamy, T. Bio-hybrid hydrogel comprising collagen-capped silver nanoparticles and melatonin for accelerated tissue regeneration in skin defects. Materials Science and Engineering: C 2021, 128. DOI: 10.1016/j.msec.2021.112328.

(19) Chen, W. H.; Chen, Q. W.; Chen, Q.; Cui, C.; Duan, S.; Kang, Y.; Liu, Y.; Liu, Y.; Muhammad, W.; Shao, S.; et al. Biomedical polymers: synthesis, properties, and applications. Sci China Chem 2022, 65 (6), 1010–1075. DOI: 10.1007/s11426-022-1243-5 From NLM PubMed-not-MEDLINE.

(20) Yousefpour, P.; McDaniel, J. R.; Prasad, V.; Ahn, L.; Li, X.; Subrahmanyan, R.; Weitzhandler, I.; Suter, S.; Chilkoti, A. Genetically Encoding Albumin Binding into Chemotherapeutic-loaded Polypeptide Nanoparticles Enhances Their Antitumor Efficacy. Nano Letters 2018, 18 (12), 7784–7793. DOI: 10.1021/acs.nanolett.8b03558.

(21) Merrett, K.; Wan, F.; Lee, C. J.; Harden, J. L. Enhanced Collagen-like Protein for Facile Biomaterial Fabrication. ACS Biomater Sci Eng 2021, 7 (4), 1414–1427. DOI: 10.1021/acsbiomaterials.1c00069 From NLM Medline.

(22) Qiu, Y.; Zhai, C.; Chen, L.; Liu, X.; Yeo, J. Current Insights on the Diverse Structures and Functions in Bacterial Collagen-like Proteins. ACS Biomater Sci Eng 2021. DOI: 10.1021/acsbiomaterials.1c00018 From NLM Publisher.

(23) Meganathan, I.; Pachaiyappan, M.; Aarthy, M.; Radhakrishnan, J.; Mukherjee, S.; Shanmugam, G.; You, J.; Ayyadurai, N. Recombinant and genetic code expanded collagen-like protein as a tailorable biomaterial. Mater Horiz 2022. DOI: 10.1039/d2mh00652a From NLM Publisher.

(24) Liu, X.; Guo, Z.; Wang, J.; Shen, W.; Jia, Z.; Jia, S.; Li, L.; Wang, J.; Wang, L.; Li, J. Thiolation-Based Protein-Protein Hydrogels for Improved Wound Healing. Advanced Healthcare Materials 2023, 2303824.

(25) Jia, S.; Wang, J.; Wang, X.; Liu, X.; Li, S.; Li, Y.; Li, J.; Wang, J.; Man, S.; Guo, Z.; et al. Genetically Encoded in situ Gelation Redox-Responsive Collagen-Like Protein Hydrogel for Accelerating Diabetic Wound Healing. Biomaterials Science 2023, 10.1039/D3BM01010D. DOI: 10.1039/D3BM01010D.

(26) Parmar, P. A.; St-Pierre, J. P.; Chow, L. W.; Puetzer, J. L.; Stoichevska, V.; Peng, Y. Y.; Werkmeister, J. A.; Ramshaw, J. A.; Stevens, M. M. Harnessing the Versatility of Bacterial Collagen to Improve the Chondrogenic Potential of Porous Collagen Scaffolds. Adv Healthc Mater 2016, 5 (13), 1656–1666. DOI: 10.1002/adhm.201600136 From NLM Medline.

(27) Bose, R. J.; Kumar, U. S.; Garcia-Marques, F.; Zeng, Y.; Habte, F.; McCarthy, J. R.; Pitteri, S.; Massoud, T. F.; Paulmurugan, R. Engineered Cell-Derived Vesicles Displaying Targeting Peptide and Functionalized with Nanocarriers for Therapeutic microRNA Delivery to Triple-Negative Breast Cancer in Mice. Advanced healthcare materials 2022, 11 (5), 2101387.

(28) Saalik, P.; Lingasamy, P.; Toome, K.; Mastandrea, I.; Rousso-Noori, L.; Tobi, A.; Simon-Gracia, L.; Hunt, H.; Paiste, P.; Kotamraju, V. R.; et al. Peptide-guided nanoparticles for glioblastoma targeting. J Control Release 2019, 308, 109–118. DOI: 10.1016/j.jconrel.2019.06.018 From NLM Medline.

(29) Hu, J.; Wang, J.; Zhu, X.; Tu, R. S.; Nanda, V.; Xu, F. Design Strategies to Tune the Structural and Mechanical Properties of Synthetic Collagen Hydrogels. Biomacromolecules 2021, 22 (8), 3440–3450. DOI: 10.1021/acs.biomac.1c00520 From NLM Medline.

(30) Squeglia, F.; Bachert, B.; De Simone, A.; Lukomski, S.; Berisio, R. The crystal structure of the streptococcal collagen-like protein 2 globular domain from invasive M3-type group A Streptococcus shows significant similarity to immunomodulatory HIV protein gp41. J Biol Chem 2014, 289 (8), 5122–5133. DOI: 10.1074/jbc.M113.523597 From NLM Medline.

(31) Wang, S.; Konorev, E. A.; Kotamraju, S.; Joseph, J.; Kalivendi, S.; Kalyanaraman, B. Doxorubicin Induces Apoptosis in Normal and Tumor Cells via Distinctly Different Mechanisms: INTERMEDIACY OF H<sub>2</sub>O<sub>2</sub>-AND p53-DEPENDENT PATHWAYS *. Journal of Biological Chemistry 2004, 279 (24), 25535–25543. DOI: 10.1074/jbc.M400944200 (acccessed 2024/12/30).

(32) Zhu, Z.; Zhai, Y.; Hao, Y.; Wang, Q.; Han, F.; Zheng, W.; Hong, J.; Cui, L.; Jin, W.; Ma, S.; et al. Specific anti-glioma targeted-delivery strategy of engineered small extracellular vesicles dual-functionalised by Angiopep-2 and TAT peptides. J Extracell Vesicles 2022, 11 (8), e12255. DOI: 10.1002/jev2.12255 From NLM Medline.

(33) Shin, J.; Jang, Y. Rational design and engineering of polypeptide/protein vesicles for advanced biological applications. Journal of Materials Chemistry B 2023.

(34) Patterson, J.; Hubbell, J. A. Enhanced proteolytic degradation of molecularly engineered PEG hydrogels in response to MMP-1 and MMP-2. Biomaterials 2010, 31 (30), 7836–7845.

(35) Liao, Z. X.; Li, Y. C.; Lu, H. M.; Sung, H. W. A genetically-encoded KillerRed protein as an intrinsically generated photosensitizer for photodynamic therapy. Biomaterials 2014, 35 (1), 500–508. DOI: 10.1016/j.biomaterials.2013.09.075 From NLM Medline.

(36) Li, X.; Yan, P.; Shao, Z. Downregulation of miR-193b contributes to enhance urokinase-type plasminogen activator (uPA) expression and tumor progression and invasion in human breast cancer. Oncogene 2009, 28 (44), 3937–3948.

(37) Sideras, K.; Braat, H.; Kwekkeboom, J.; Van Eijck, C.; Peppelenbosch, M.; Sleijfer, S.; Bruno, M. Role of the immune system in pancreatic cancer progression and immune modulating treatment strategies. Cancer treatment reviews 2014, 40 (4), 513–522.

(38) Abbas, M.; Zou, Q.; Li, S.; Yan, X. Self-assembled peptide-and protein-based nanomaterials for antitumor photodynamic and photothermal therapy. Advanced materials 2017, 29 (12), 1605021.

(39) Sheng, S.; Zhang, H.; Li, X.; Chen, J.; Wang, P.; Liang, Y.; Li, C.; Li, H.; Pan, N.; Bao, X. Probiotic-derived amphiphilic exopolysaccharide self-assembling adjuvant delivery platform for enhancing immune responses. Journal of Nanobiotechnology 2024, 22 (1), 267.

(40) Wang, J.; Li, J.; Sun, Y.; Liu, X.; Wang, L.; Xia, Y.; Huang, J.; Feng, J.; Jia, S.; Li, Y.; et al. Genetically Encoded Incorporation of IFN-α into Collagen-like Protein–Hyaluronic Acid Hydrogels for Diabetic Chronic Wound Healing. ACS Materials Letters 2024, 4133–4141. DOI: 10.1021/acsmaterialslett.4c01170.

