## Supplementary for "Chimeric Collagen-like Proteins with Tunable Structural Heterogeneity for Precise Control and Targeted Doxorubicin Delivery"

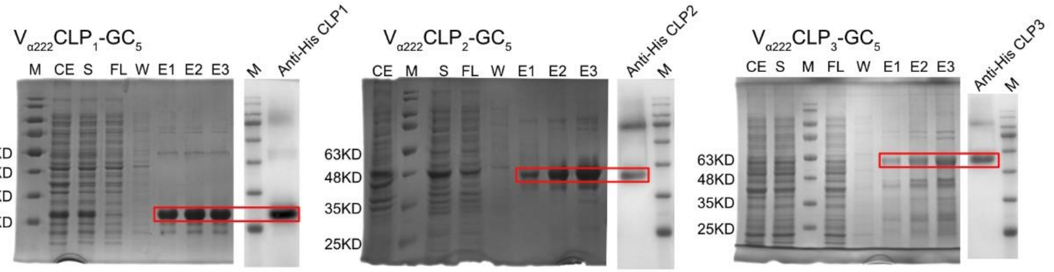


Figure. S1: Protein purification SDS-PAGE and WB validation of CLPs with different linear structure lengths.

| | $V_{\alpha222}$CLP1-GC$_5$-Dox | $V_{\alpha222}$CLP2-GC$_5$-Dox | $V_{\alpha222}$CLP3-GC$_5$-Dox |
|---|---|---|---|
| **Zeta Potential (mV)** | -11.5 | -13.8 | -22.7 |

Table. S1: Zeta potentials of CDCNs made of linear structural protein CLPs of different lengths.

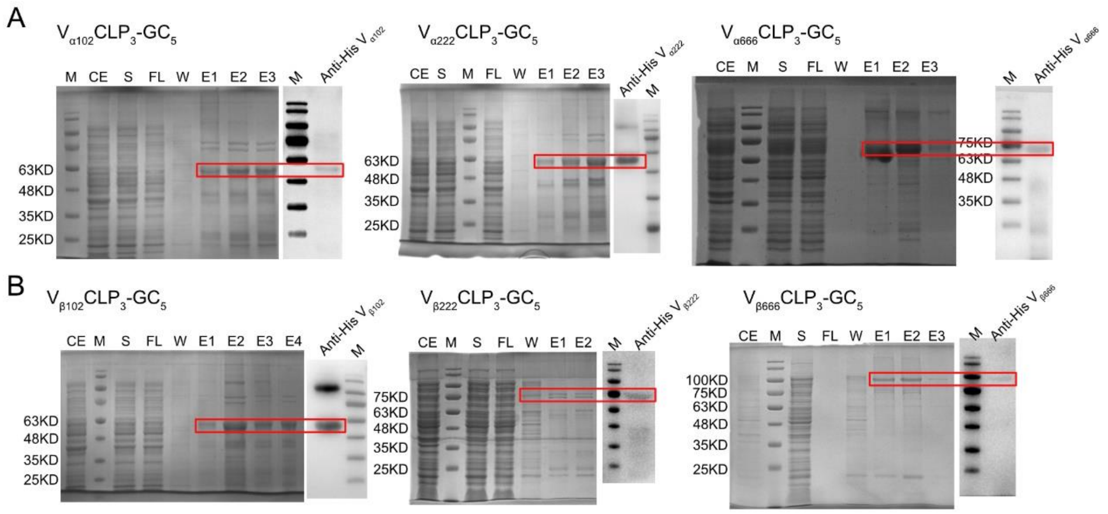


Figure. S2: Protein purification SDS-PAGE and WB validation of the proteins with N-terminal α-helix (A) and β-sheet (B) spherical structural domains of different lengths.

| | $V_{\alpha102}CLP_3$-$GC_5$-Dox | $V_{\alpha222}CLP_3$-$GC_5$-Dox | $V_{\alpha666}CLP_3$-$GC_5$-Dox |
|---|---|---|---|
| **Zeta Potential (mV)** | -25.3 | -23.8 | -21.3 |
| | **$V_{\beta102}CLP_3$-$GC_5$-Dox** | **$V_{\beta222}CLP_3$-$GC_5$-Dox** | **$V_{\beta666}CLP_3$-$GC_5$-Dox** |
| **Zeta Potential (mV)** | -16.8 | -14.7 | -13.9 |

Table. S2: Zeta potentials of CDCNs made of proteins with N-terminal α-helix and β-sheet spherical structural domains of different lengths.

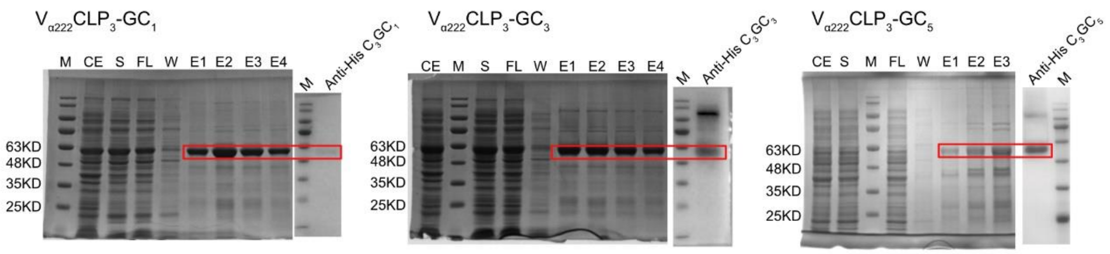


Figure. S3: Protein purification SDS-PAGE and WB validation of the purification of proteins CLPs with varying DOX conjugation sites.

| | $V_{\alpha222}CLP_3$-GC1-Dox | $V_{\alpha222}CLP_3$-GC3-Dox | $V_{\alpha222}CLP_3$-GC5-Dox |
|---|---|---|---|
| **Zeta Potential (mV)** | -16.1 | -18.4 | -22.7 |

Table. S3: Zeta potentials of CDCNs made of proteins CLPs with varying DOX conjugation sites.

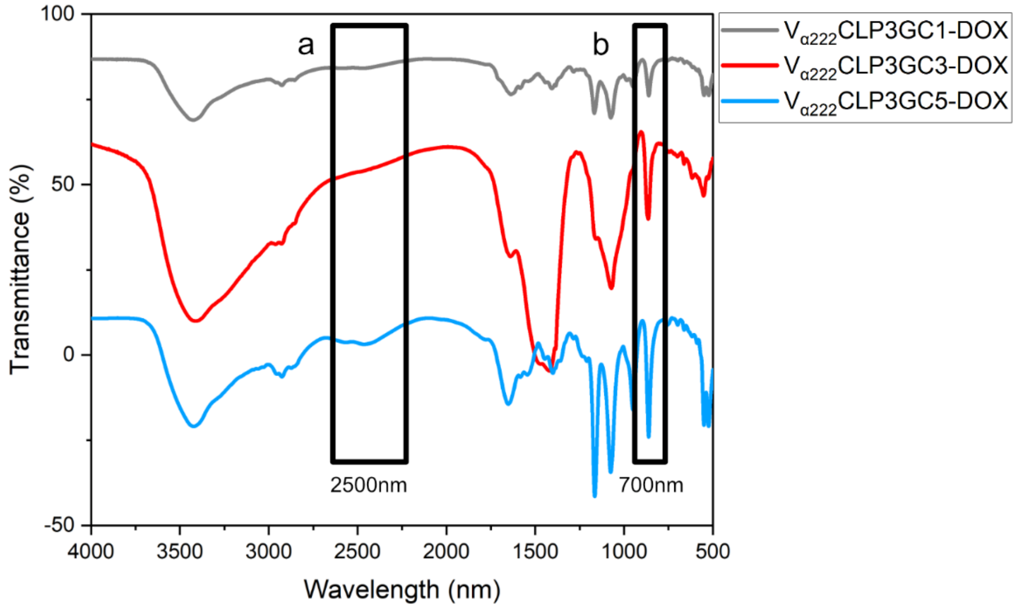


Figure. S4: FTIR spectra of CDCNs at DOX drug attachment points 1, 3, and 5. The a-peak is the S-H vibration of cysteine, and the b-peak is the thioether bonding vibration.

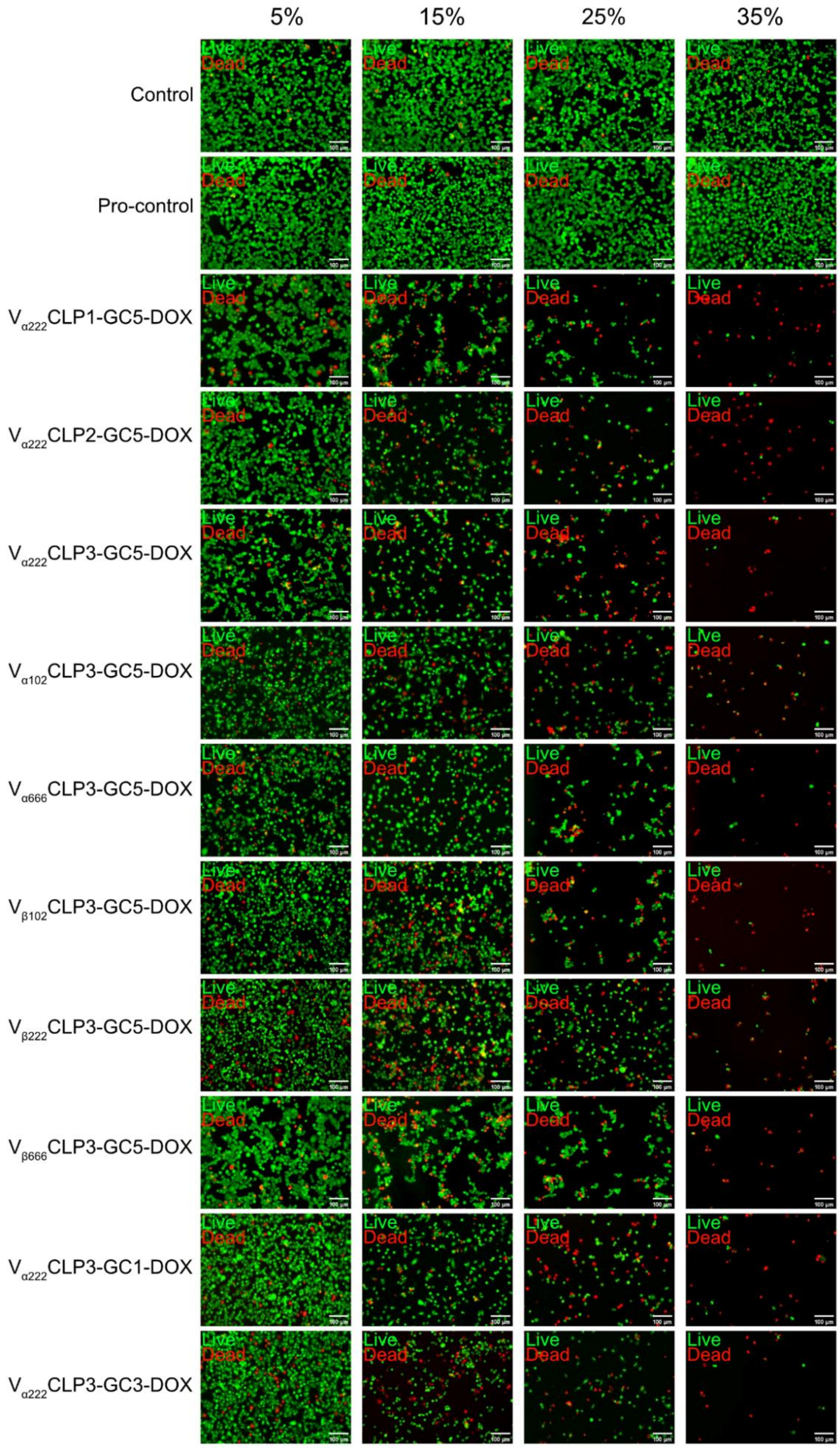


Figure. S5: Results of dead-viable staining of 4T1 cells by CDCNs at different concentration gradients.

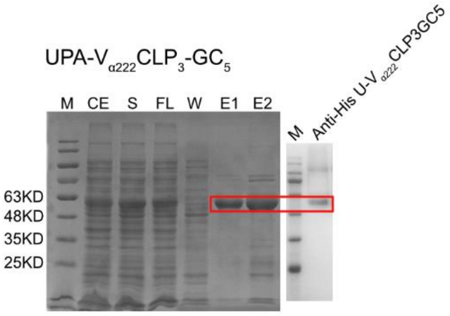


Figure. S6: Protein purification SDS-PAGE and WB validation of the purification of targeting peptides UPA added to the N-terminus of $V_{\alpha222}$CLP3GC5.

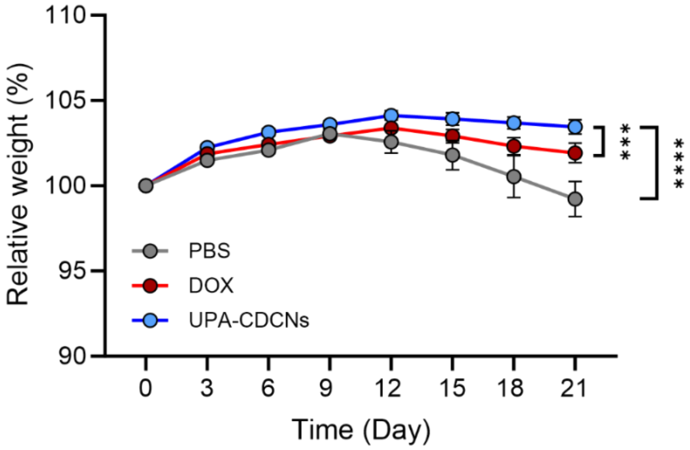


Figure. S7: Relative body weight curves in TNBC mice.

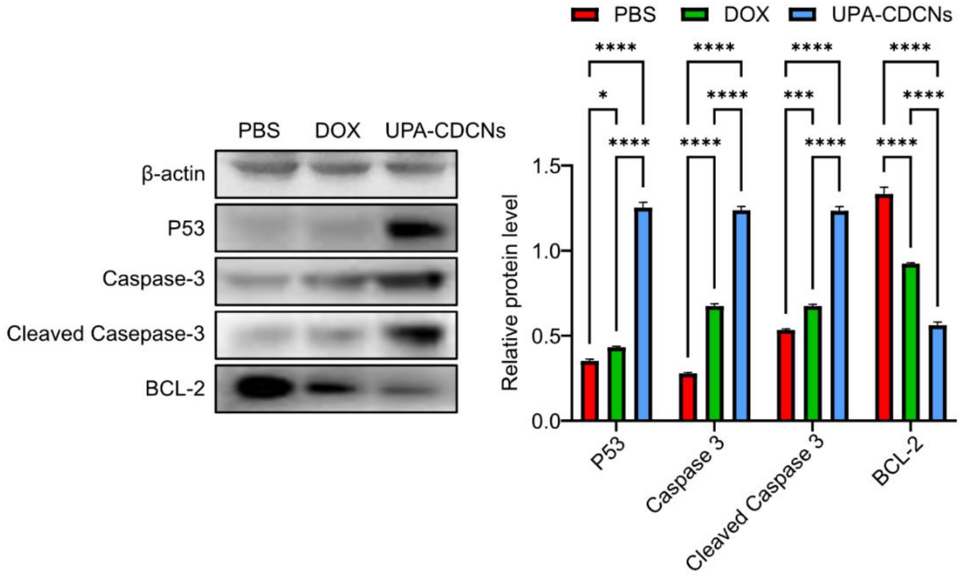


Figure. S8: WB validation of three groups of breast cancer tumors with PBS, DOX, UPA-CDCNs and their grayscale analysis.

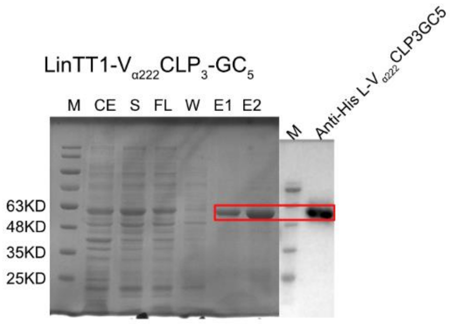


Figure. S9: Protein purification SDS-PAGE and WB validation of the purification of targeting peptides LinTT1 added to the N-terminus of $V_{\alpha222}$CLP3GC5.

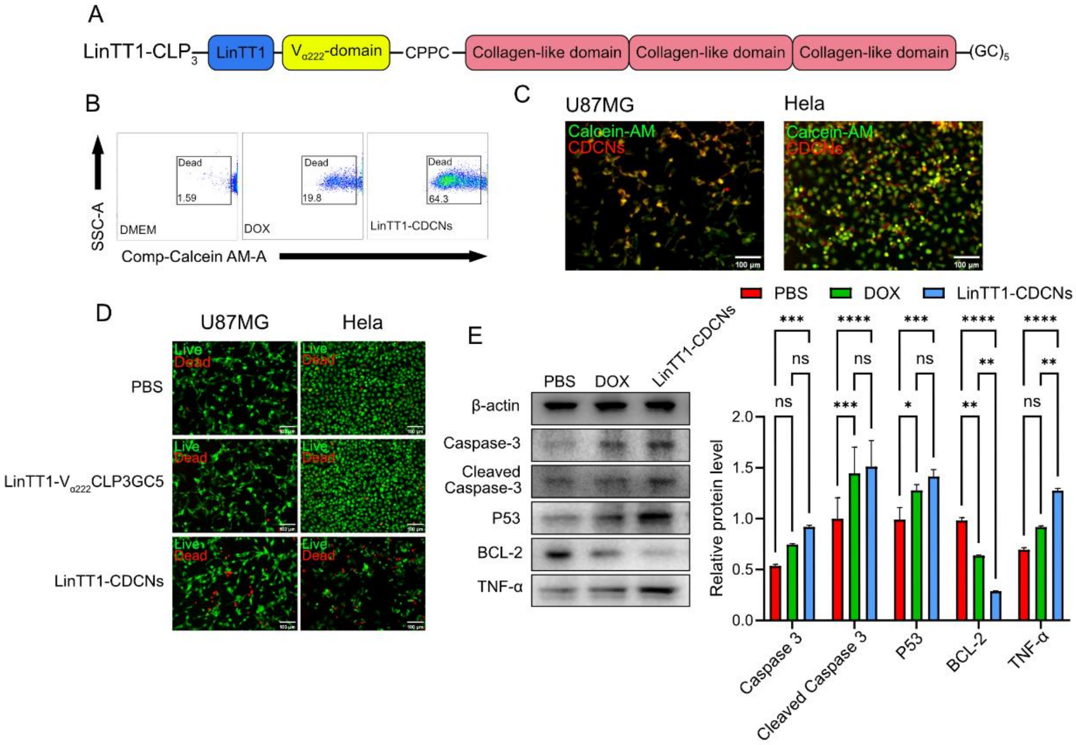


Figure. S10: Effects of LinTT1-CDCNs on GBM cells.

(A). Schematic representation of the construction of GBM-targeting CLPs. The GBM-targeting peptide, LinTT1, was genetically encoded at the N-terminal of $V_{\alpha222}$CLP3GC5 to generate LinTT1-$V_{\alpha222}$CLP3GC5.

(B). Mortality was examined using flow cytometry on DMEM control, DOX and LinTT1-CDCNs.

(C). Imaging of LinTT1-CDCNs targeting ability. Cells were stained with Calcein AM and imaged in the FITC channel. Red fluorescence represents the spontaneous fluorescence of DOX, captured in the TRITC channel.

(D). Live/dead staining images to assess the anti-tumor effects of LinTT1-CDCNs.

(E). Western blot analysis and quantitative statistics of 4T1 cells after treatment with PBS, DOX, and LinTT1-CDCNs.

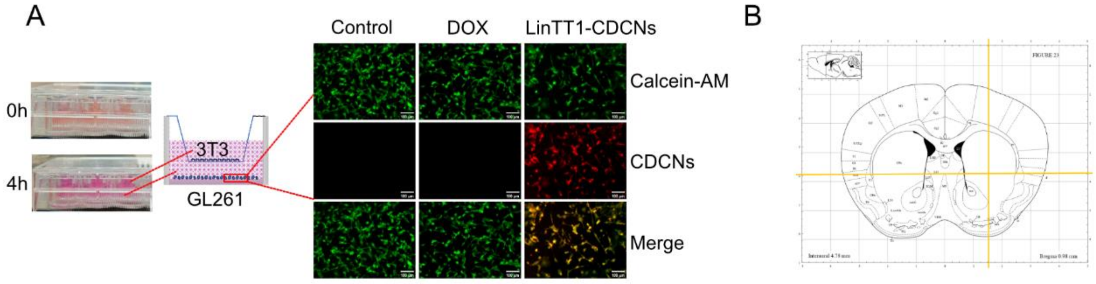


Figure. S11: (A) In vitro permeabilization assay to verify the ability of LinTT1-CDCNs to cross the blood-brain barrier. (B) Mouse stereotaxic brain localization injection sites.

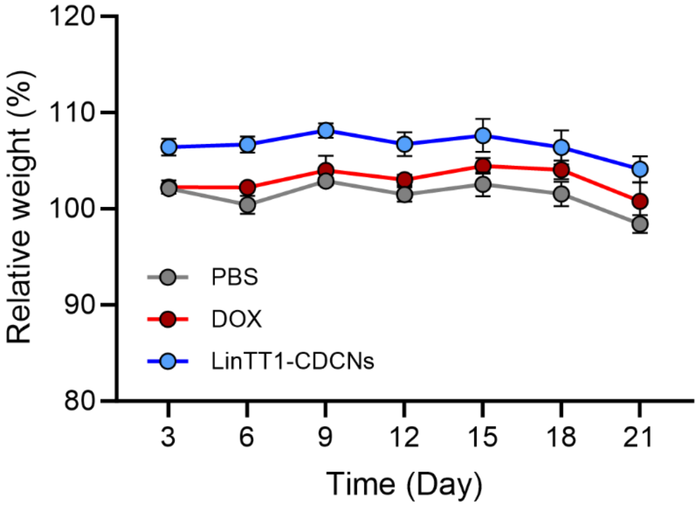


Figure. S12: Relative body weight curves in GBM mice.

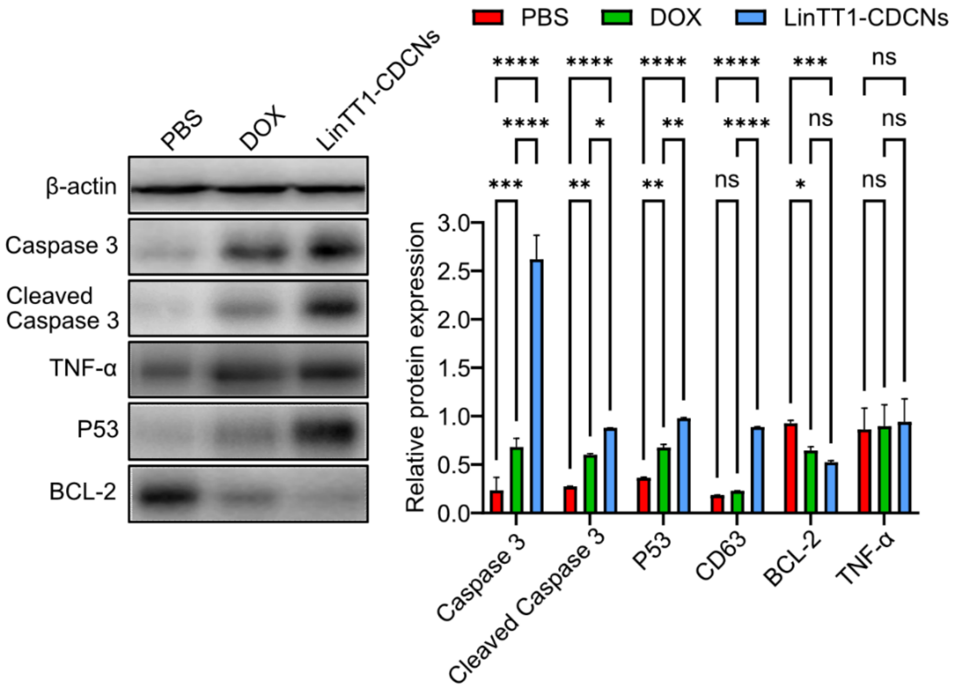


Figure. S13: WB validation of three groups of brain protein in mice with glioma tumors with PBS, DOX, LinTT1-CDCNs and their grayscale analysis.

| Name of gene(s) | **Amino acid sequences** |
| --- | --- |
| $V_{\alpha222}$CLP1-GC5 | MADEQEEKAKVRTELIQELAQGLGGIEKKNFPTLGDEDLDHTYMTKL LTYLQEREQAENSWRKRLLKGIQDHALDGGPCPPCGPKGEQGPQGLP GKDGEAGAQGPAGPMGPAGEQGEKGEPGTQGAKEDRGETGPKGPK GERGEAGPAGKDGEPGPVGPARGDGCGCGCGCGC |
| $V_{\alpha222}$CLP2-GC5 | MADEQEEKAKVRTELIQELAQGLGGIEKKNFPTLGDEDLDHTYMTKL LTYLQEREQAENSWRKRLLKGIQDHALDGGPCPPCGPKGEQGPQGLP GKDGEAGAQGPAGPMGPAGEQGEKGEPGTQGAKEDRGETGPKGPK GERGEAGPAGKDGEPGPVGPAGPKGEQGPQGLPGKDGEAGAQGPAG PMGPAGEQGEKGEPGTQGAKEDRGETGPKGPKGERGEAGPAGKDGE PGPVGPARGDGCGCGCGCGC |
| $V_{\alpha222}$CLP3-GC5 | MADEQEEKAKVRTELIQELAQGLGGIEKKNFPTLGDEDLDHTYMTKL LTYLQEREQAENSWRKRLLKGIQDHALDGGPCPPCGPKGEQGPQGLP GKDGEAGAQGPAGPMGPAGEQGEKGEPGTQGAKEDRGETGPKGPK GERGEAGPAGKDGEPGPVGPAGPKGEQGPQGLPGKDGEAGAQGPAG PMGPAGEQGEKGEPGTQGAKEDRGETGPKGPKGERGEAGPAGKDGE PGPVGPAGPKGEQGPQGLPGKDGEAGAQGPAGPMGPAGEQGEKGEP GTQGAKEDRGETGPKGPKGERGEAGPAGKDGEPGPVGPARGDGCGC GCGCGC |
| $V_{\alpha102}$CLP3-GC5 | MSIPITKAQLRTFRAIIIDLTKIIPKLFANPSPQNGGPCPPCGPKGEQGPQ GLPGKDGEAGAQGPAGPMGPAGEQGEKGEPGTQGAKEDRGETGPKG PKGERGEAGPAGKDGEPGPVGPAGPKGEQGPQGLPGKDGEAGAQGP AGPMGPAGEQGEKGEPGTQGAKEDRGETGPKGPKGERGEAGPAGKD GEPGPVGPAGPKGEQGPQGLPGKDGEAGAQGPAGPMGPAGEQGEKG EPGTQGAKEDRGETGPKGPKGERGEAGPAGKDGEPGPVGPARGDGC GCGCGCGC |
| $V_{\alpha666}$CLP3-GC5 | MSIPITKAQLRTFRAIIIDLTKIIPKLFANPSPQNIEDLIDTLNLLSKFICSL EATSSLKAQGLAIIKNLITILKNPTFVASAVFVELQILINYLLYITKLFRI DHCTLQELSIPITKAQLRTFRAIIIDLTKIIPKLFANPSPQNIEDLIDTLNL LSKFICSLEATSSLKAQGLAIIKNLITILKNPTFVASAVFVELQILINYLL YITKLFRIDHCTLQELGGPCPPCGPKGEQGPQGLPGKDGEAGAQGPAG PMGPAGEQGEKGEPGTQGAKEDRGETGPKGPKGERGEAGPAGKDGE PGPVGPAGPKGEQGPQGLPGKDGEAGAQGPAGPMGPAGEQGEKGEP GTQGAKEDRGETGPKGPKGERGEAGPAGKDGEPGPVGPAGPKGEQG PQGLPGKDGEAGAQGPAGPMGPAGEQGEKGEPGTQGAKEDRGETGP KGPKGERGEAGPAGKDGEPGPVGPARGDGCGCGCGCGC |

| | |
|---|---|
| $V_{\beta102}$CLP3-GC5 | MHQGSGGSQGTAAGETGQSDIPGSAGSQGTIGNETGGPCPPCGPKGE<br>QGPQGLPGKDGEAGAQGPAGPMGPAGEQGEKGEPGTQGAKEDRGET<br>GPKGPKGERGEAGPAGKDGEPGPVGPAGPKGEQGPQGLPGKDGEAG<br>AQGPAGPMGPAGEQGEKGEPGTQGAKEDRGETGPKGPKGERGEAGP<br>AGKDGEPGPVGPAGPKGEQGPQGLPGKDGEAGAQGPAGPMGPAGEQ<br>GEKGEPGTQGAKEDRGETGPKGPKGERGEAGPAGKDGEPGPVGPAR<br>GDGCGCGCGCGC |
| $V_{\beta222}$CLP3-GC5 | MSGWSTATTGDTYGVWGLCDSNAGVGVVGHARATTGLTTGVLGRA<br>DAASGDAAGVYGYASGATGATAGVWGVVESGGPCPPCGPKGEQGP<br>QGLPGKDGEAGAQGPAGPMGPAGEQGEKGEPGTQGAKEDRGETGPK<br>GPKGERGEAGPAGKDGEPGPVGPAGPKGEQGPQGLPGKDGEAGAQG<br>PAGPMGPAGEQGEKGEPGTQGAKEDRGETGPKGPKGERGEAGPAGK<br>DGEPGPVGPAGPKGEQGPQGLPGKDGEAGAQGPAGPMGPAGEQGEK<br>GEPGTQGAKEDRGETGPKGPKGERGEAGPAGKDGEPGPVGPARGDG<br>CGCGCGCGC |
| $V_{\beta666}$CLP3-GC5 | MGQSDIQGSTGSQGTAGGQETGQSDIQGSSGSQGTAGNESGISDHQGSG<br>GSQGTAAGETGQSDIPGSAGSQGTIGNETGQSDIQGSTGSQGTAGQET<br>GQSDIQGSTGSQGTAGAETGQSDVQGSGGSVGTAGAETGSSDVQGST<br>GSAGTAGAETGSSDVQGSTGSQGTAGAETGHSDVQGSTGSQGTAGN<br>ESGISDVQGSTGSQGTAGNETGGSDVQGSTGSVGGGPCPPCGPKGEQ<br>GPQGLPGKDGEAGAQGPAGPMGPAGEQGEKGEPGTQGAKEDRGETG<br>PKGPKGERGEAGPAGKDGEPGPVGPAGPKGEQGPQGLPGKDGEAGA<br>QGPAGPMGPAGEQGEKGEPGTQGAKEDRGETGPKGPKGERGEAGPA<br>GKDGEPGPVGPAGPKGEQGPQGLPGKDGEAGAQGPAGPMGPAGEQG<br>EKGEPGTQGAKEDRGETGPKGPKGERGEAGPAGKDGEPGPVGPARG<br>DGCGCGCGCGC |
| $V_{\alpha222}$CLP3-GC1 | MADEQEEKAKVRTELIQELAQGLGGIEKKNFPTLGDEDLDHTYMTKL<br>LTYLQEREQAENSWRKRLLKGIQDHALDGGPCPPCGPKGEQGPQGLP<br>GKDGEAGAQGPAGPMGPAGEQGEKGEPGTQGAKEDRGETGPKGPK<br>GERGEAGPAGKDGEPGPVGPAGPKGEQGPQGLPGKDGEAGAQGPAG<br>PMGPAGEQGEKGEPGTQGAKEDRGETGPKGPKGERGEAGPAGKDGE<br>PGPVGPAGPKGEQGPQGLPGKDGEAGAQGPAGPMGPAGEQGEKGEP<br>GTQGAKEDRGETGPKGPKGERGEAGPAGKDGEPGPVGPARGDGC |
| $V_{\alpha222}$CLP2-GC3 | MADEQEEKAKVRTELIQELAQGLGGIEKKNFPTLGDEDLDHTYMTKL<br>LTYLQEREQAENSWRKRLLKGIQDHALDGGPCPPCGPKGEQGPQGLP<br>GKDGEAGAQGPAGPMGPAGEQGEKGEPGTQGAKEDRGETGPKGPK<br>GERGEAGPAGKDGEPGPVGPAGPKGEQGPQGLPGKDGEAGAQGPAG<br>PMGPAGEQGEKGEPGTQGAKEDRGETGPKGPKGERGEAGPAGKDGE<br>PGPVGPAGPKGEQGPQGLPGKDGEAGAQGPAGPMGPAGEQGEKGEP<br>GTQGAKEDRGETGPKGPKGERGEAGPAGKDGEPGPVGPARGDGCGC<br>GC |

| | |
|---|---|
| UPA-$V_{\alpha222}$CLP3-GC5 | MVSNKYFSNIHWGCADEQEEKAKVRTELIQELAQGLGGIEKKNFPTLGDEDLDHTYMTKLLTYLQEREQAENSWRKRLLKGIQDHALDGGPCPPCGPKGEQGPQGLPGKDGEAGAQGPAGPMGPAGEQGEKGEPGTQGAKEDRGETGPKGPKGERGEAGPAGKDGEPGPVGPAGPKGEQGPQGLPGKDGEAGAQGPAGPMGPAGEQGEKGEPGTQGAKEDRGETGPKGPKGERGEAGPAGKDGEPGPVGPAGPKGEQGPQGLPGKDGEAGAQGPAGPMGPAGEQGEKGEPGTQGAKEDRGETGPKGPKGERGEAGPAGKDGEPGPVGPARGDGCGCGCGCGC |
| LinTT1-$V_{\alpha222}$CLP3-GC5 | MAKRGARSTAADEQEEKAKVRTELIQELAQGLGGIEKKNFPTLGDEDLDHTYMTKLLTYLQEREQAENSWRKRLLKGIQDHALDGGPCPPCGPKGEQGPQGLPGKDGEAGAQGPAGPMGPAGEQGEKGEPGTQGAKEDRGETGPKGPKGERGEAGPAGKDGEPGPVGPAGPKGEQGPQGLPGKDGEAGAQGPAGPMGPAGEQGEKGEPGTQGAKEDRGETGPKGPKGERGEAGPAGKDGEPGPVGPAGPKGEQGPQGLPGKDGEAGAQGPAGPMGPAGEQGEKGEPGTQGAKEDRGETGPKGPKGERGEAGPAGKDGEPGPVGPARGDGCGCGCGCGC |

Table. S4: Amino acid sequences of genes used in the article.
